# Identification of the targets of T cell receptor therapeutic agents and cells by use of a high throughput genetic platform

**DOI:** 10.1101/267047

**Authors:** Ron S. Gejman, Heather F. Jones, Martin G. Klatt, Aaron Y. Chang, Claire Y. Oh, Smita S. Chandran, Tatiana Korontsvit, Viktoriya Zakahleva, Tao Dao, Christopher A. Klebanoff, David A. Scheinberg

**Affiliations:** Molecular Pharmacology Program, Memorial Sloan Kettering Cancer Center, New York, NY, USA. 10065; Tri-Institutional MD-PhD Program (MSKCC, Rockefeller University, Weill Cornell Medical College), New York, NY, USA. 10065; Weill Cornell Medicine, New York, NY, USA. 10065; Center for Cell Engineering and Department of Medicine, MSKCC, New York, NY, USA. 10065; Parker Institute for Cancer Immunotherapy, MSKCC, New York, NY, USA. 10065

## Abstract

T cell receptor (TCR)-based therapeutic cells and agents have emerged as a new class of effective cancer therapeutics. These therapies work on cells that express intracellular cancer-associated proteins by targeting peptides displayed on major histocompatibility complex receptors. However, cross-reactivities of these agents to off-target cells and tissues have resulted in serious, sometimes fatal, adverse events. We have developed a high throughput genetic platform (termed “PresentER”) that encodes MHC-I peptide minigenes for functional immunological assays as well as for determining the reactivities of TCR-like therapeutic agents against large libraries of MHC-I ligands. In this report, we demonstrate that PresentER can be used to identify the on-and-off targets of T cells and TCR mimic antibodies using *in vitro* co-culture assays or binding assays. We find dozens of MHC-I ligands that are cross-reactive with two TCR mimic antibodies and two native TCRs and that are not easily predictable by other methods.

## Introduction

T cell receptor (TCR)-based therapeutic cells and agents — including adoptive T cells and tumor infiltrating lymphocytes (TILs)^1,2^, TCR-engineered T cells^3^, ImmTACs^4^, TCR mimic antibodies^5^, and neoantigen vaccines^6,7^—are cancer therapeutics that can target cells expressing intracellular cancer-associated proteins. These agents rely on presentation of short peptides derived from cellular, viral or phagocytosed proteins on major histocompatibility complex (MHC), also known as human leukocyte antigen (HLA). However, cross-reactivities of these agents to off-target cells and tissues are poorly understood, difficult to predict, and have resulted in serious, sometimes fatal, adverse events^8,9^. In addition, identifying the antigenic targets of therapeutic TIL’s found in tumors is time-consuming, expensive, and complicated^2^.

Here we have developed a mammalian minigene-based system (termed “PresentER”) that encodes libraries of MHC-I peptide ligands for determining the reactivities and cross-reactivities of TCR-like therapeutic drugs. PresentER encoded minigenes produce short (8-11aa) peptides that are translated directly into the endoplasmic reticulum using a signal sequence and thus bypass the endogenous protein processing steps that produce MHC-I peptides from full-length proteins. For the purpose of identifying the targets of T cells, the approach described herein is superior to heterologous expression of full-length cDNA as it avoids the unpredictable effects of peptide processing (including proteasome cleavage, transporter associated with antigen processing (TAP) and N-terminal trimming by aminopeptidases). Here we show that genetically encoded MHC-I peptides can be used in biochemical and live-cell cytotoxicity assays to identify the on-target and off-target ligands of soluble TCR multimers, TCR mimic (TCRm) antibodies and engineered T cell receptors expressed on lymphocytes. PresentER expressing cells can also be used in immunologic assays such as T cell activation, cytotoxicity and tumor rejection in naïve, wild-type mice^10^. Using PresentER in pooled library screens, we are able to rapidly discover dozens of peptide ligands of two soluble TCR mimic antibodies and two engineered-TCR T cells, from among thousands of potential epitopes.

TCR based therapeutics are structurally similar to the TCR on CD8 T cells and thus share both their potential advantages and challenges. For instance, CD8 T cells can theoretically discern whether any MHC-I bound peptide is self, foreign or altered-self. Yet, the number of possible MHC-I ligands that can be encoded by the twenty proteinogenic amino acids is significantly larger than the number of circulating T cells in the human body. In order to account for this discrepancy, TCR are highly cross-reactive: a single TCR may be capable of recognizing over 1 million distinct pMHC^11^. Thymic selection *in vivo* is critical to reduce circulating T cells that may be auto-reactive. Some TCR-based therapeutics do not undergo negative selection for the human pMHC repertoire or, if they are isolated from humans, they are sometimes modified to make them higher affinity. As a consequence, each of these agents can be cross-reactive with HLA presented peptides found in normal tissue^12^. A prominent example is an affinity-enhanced TCR directed against an HLA-A*01:01 MAGE-A3 peptide (168-176; EVDPIGHLY), which induced lethal cardiotoxicity when two patients were treated with these T cells during a phase I clinical trial. Extensive preclinical testing failed to uncover off-target reactivity, but afterwards it was discovered that an epitope derived from Titin (24337-24345; ESDPIVAQY), a structural protein highly expressed by cardiomyocytes, was cross-reactive with the MAGE-A3 TCR^8^. Another TCR directed towards the MAGE-A3 peptide (112-120: KVAELVHFL) led to neuronal toxicity and death in several treated patients, thought to be due to cross-reactivity of the TCR to a peptide from the MAGE-A12 protein (112-120: KMAELVHFL)^9^. Hence, a major challenge to the development of safe TCR based therapeutics is the prospective identification of off-tumor, off-target pMHC^13^.

Identifying off-tumor, off-target pMHC is challenging for two main reasons: (1) the scope of the problem is undefined because the complete repertoire of HLA ligands found in normal human tissue is unknown. The number of known HLA ligands in humans is rapidly expanding, with recent reports identifying thousands of novel presented peptides^14,15^. However, it is unclear how many presented peptides remain to be discovered and little is known about the antigens presented on important tissues such as the nerves, eyes, heart and lungs. (2) Cross-reactive pMHC are not easy to identify. Methods to identify cross-reactive targets of TCR-like molecules have been developed by panning yeast^16,17^ or insect-baculovirus^18^ cells against soluble TCRs or by staining T cells with libraries of pMHC-tetramers^19,20^. These approaches are highly valuable, but each has caveats, including time-consuming bacterial purification/refolding of soluble TCRs or expensive synthesis of peptides to make tetramers. Finally, these systems do not mimic the most important aspect of T cell therapy: that is, killing of a target cell. Here, we show that the PresentER system is physiologically relevant, scaleable, sensitive, specific and can rapidly identify functional cross-reactivities between MHC-I ligands and TCR-like agents in biochemical binding assays, immunological assays *in vitro* and *in vivo*, such as T cell cytotoxicity assays and tumor rejection. For each of the 4 TCR agents studied in this manuscript, multiple cross-reactive peptides were identified, including some which are expressed and presented on human cells.

This approach offers two specific advantages over other approaches to generate MHC-I that have been used in the field: (1) avoidance of unpredictable endogenous peptide processing steps resulting in MHC-I molecules bearing unknown peptides and (2) peptides presented on MHC are bound non-covalently, as actually occurs in real cells—as opposed to using a flexible linker covalently tethered to an engineered MHC molecule.

## Results

### PresentER yield functional MHC-I ligands for biochemical and functional immunological assays

We designed a minigene (“PresentER”) that is capable of generating precisely defined MHC-I antigen in mammalian cells. The DNA encoding the MHC-I peptide is short (72-78nt) and inexpensive to synthesize individually or as a pool (**Figure 1A and Fig. S1)**. The peptide is encoded downstream of a signal sequence, thus bypassing the typical processing for MHC-I peptide presentation: proteasomal MHC-I H-2Kb/SIINFEKL bound the correct epitope in the Tap2 deficient mouse RMA/S cell line expressing PresentER SIINFEKL (Chicken Ovalbumin 257-264) versus irrelevant minigene MSIIFFLPL (PEDF:271–279) (**Fig. 1C)**. Cell lines with wild-type Tap function could also yield PresentER driven antigens, as shown by the expression of PresentER-SIINFEKL in EL4 cells **(Fig. S3)**.

**Fig. 1:**
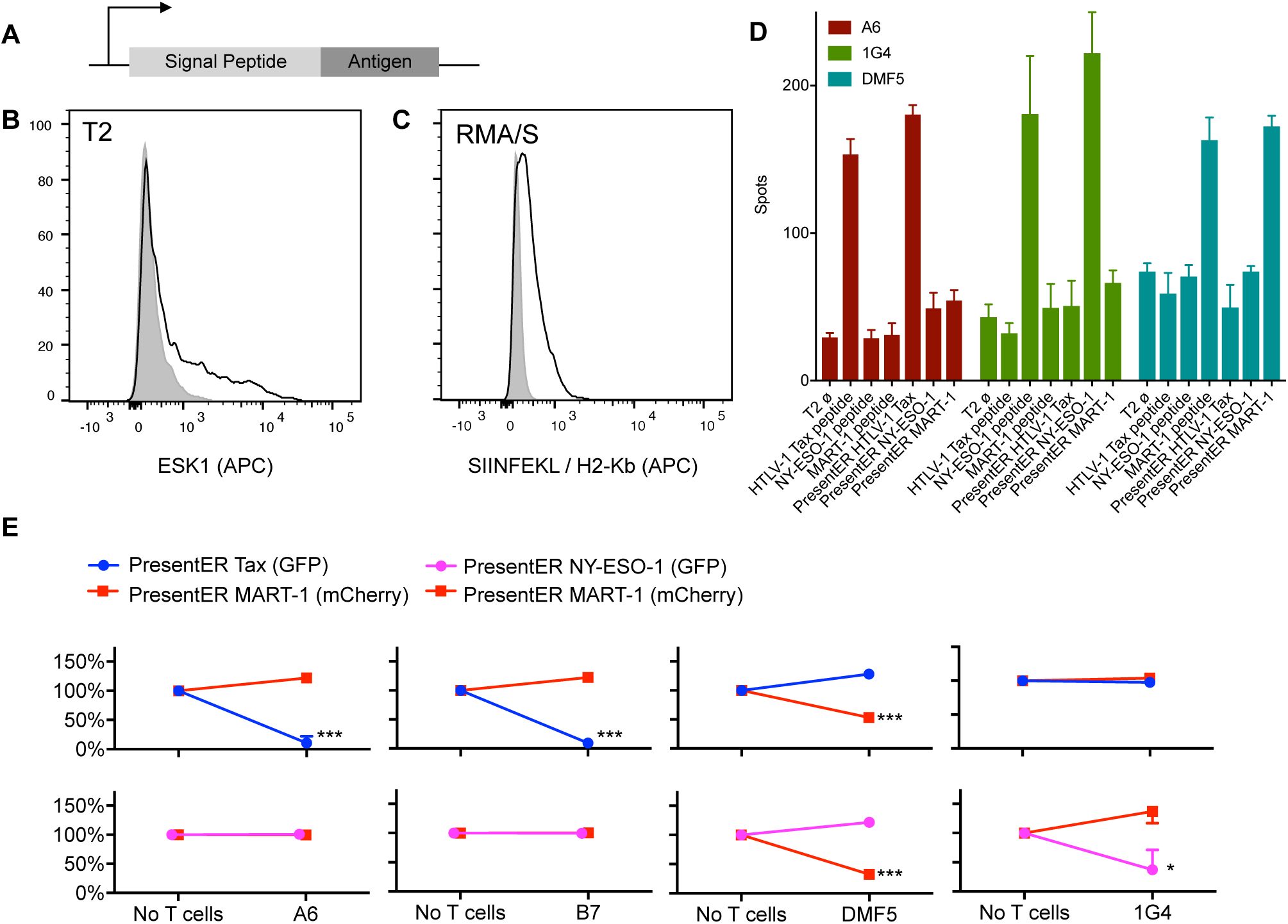
Design and characterization of PresentER minigene. **(A)** The PresentER minigene encodes an ER signal sequence, followed by a peptide antigen and a stop codon. **(B)** T2 cells expressing PresentER-RMFPNAPYL (black) but not PresentER-ALYVDSLFFL (gray) are bound by fluorescently labeled ESK1, a TCR mimic antibody to the complex of RMFPNAPYL/HLA-A*02:01. **(C)** A fluorescently labeled antibody to SIINFEKL/H-2Kb binds to RMA/S cells expressing PresentER-SIINFEKL (black), but not to PresentER-MSIIFFLPL (gray). **(D)** ELISPOT of genetically engineered T cells expressing the A6 (target: HTLV-1 Tax:11-19 LLFGYPVYV), DMF5 (target: MART-1:27-35 AAGIGILTV) or 1G4 (target: NY-ESO-1:157-165 SLLMWITQC) TCRs challenged with peptide-pulsed T2 cells or T2 cells expressing PresentER minigenes. **(E)** Results of *in vitro* co-culture killing assays where A6, B7, DMF5 and 1G4 expressing T cells were incubated with a mixture of T2 cells expressing PresentER-Tax (GFP)/PresentER MART-1 (mCherry), PresentER NY-ESO-1 (GFP)/PresentER MART-1 (mCherry). The change in abundance of the T2 target cells are plotted relative to their abundance in the “No T cells” sample.

We immunoprecipitated peptide-MHC complexes from T2 cells expressing PresentER-RMFPNAPYL or PresentER-cleavage, peptide transport into the endoplasmic reticulum and aminopeptidase trimming^21^. The peptide is translated directly into the endoplasmic reticulum, where it binds to MHC and is exported to the surface of the cell. PresentER was designed for pooled screening applications; therefore, the amino acid sequence corresponding to a peptide is encoded by DNA only once per minigene and can be sequenced in its entirety in one short (50bp) next generation sequencing read. In order to first demonstrate that PresentER minigenes recapitulate all of the expected characteristics of MHC-I presented antigens, we relied on several fluorescently labeled monoclonal antibodies, multimerized TCRs and engineered T cells. All TCR and TCR-like agents used, together with their target peptides, are described in **Table 1**. These agents serve as reporters to indicate that mature MHC-I molecules bearing the encoded peptides are indeed being generated by the minigene.

**Table 1:**
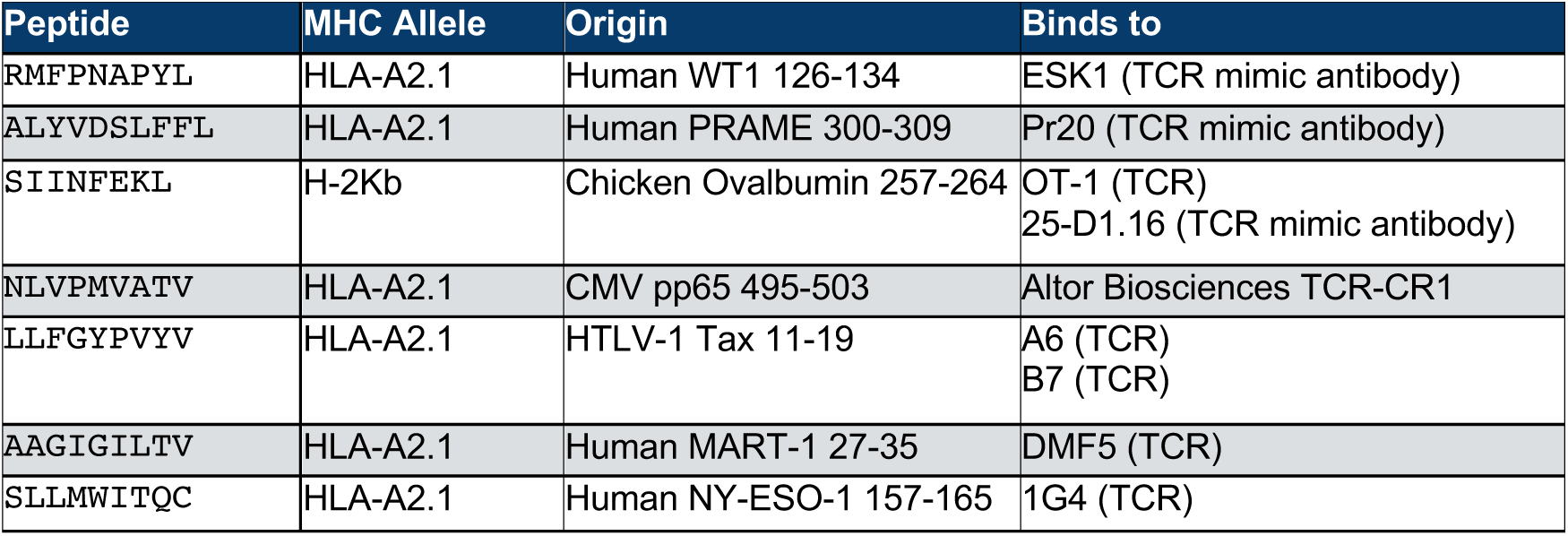
The target of each TCR and TCR-like agent used in this article.

We have previously isolated and characterized two TCR mimic (TCRm) antibodies (ESK1^5^ and Pr20^22^) that recognize HLA-A*02:01 (HLA-A2.1) peptides derived from genes aberrantly expressed on cancer cells. ESK1 binds to the Wilms tumor (WT1) peptide RMFPNAPYL (WT1:126-134) and Pr20 binds to the Preferentially expressed antigen of melanoma (PRAME) peptide ALYVDSLFFL (PRAME:300-309). When T2 cell lines expressing a variety of PresentER-encoded peptides were stained with fluorescently labeled ESK1 and Pr20, these antibodies bound only cells expressing their cognate antigens, but not irrelevant peptides **(Fig. 1B and Fig. S2A-D)**. In order to show that TCR could bind to the minigene-derived MHC-I peptides, we used two HLA-A2.1 peptide specific TCR multimers: (1) TCR-CR1 which recognizes NLVPMVATV (Cytomegalovirus pp65:495-503) and (2) A6 which recognizes LLFGYPVYV (HTLV-1 Tax:11-19)^23^. These two soluble TCR multimers specifically bound T2 cells expressing their cognate ligand **(Fig. S2E-H).**

PresentER encoded minigenes could also yield MHC-I peptides to other MHC alleles: an antibody against mouse ALYVDSLFFL and identified bound peptides by mass spectrometry. RMFPNAPYL and ALYVDSLFFL were identified only in cells encoding those PresentER constructs (**Fig. S4A-B**). No peptides derived from the ER signal sequence were identified. We scrambled the ER signal sequence to test whether peptides were associating with MHC via a signal-sequence independent mechanism and found no binding to cells expressing these constructs **(Fig. S4C)**. Finally, by transducing cells with serially diluted viral supernatant, thus achieving low multiplicity of infection (MOI), we show that the PresentER minigene is capable of driving antigen presentation from a single copy of the retroviral minigene, thereby enabling its use in a pooled screen **(Fig. S5)**.

Next, we wanted to test if PresentER encoded MHC-I antigens could be used in functional immunology assays using T cells. Genetically engineered T cells expressing the DMF5^3,24^ (specific to MART-1:27-35 HLA-A2.1/AAGIGILTV), 1G4^25^ (specific to NY-ESO-1:157-165 HLA-A2.1/SLLMWITQC) and A6 TCRs released IFN-γ when exposed to peptide-pulsed cells or cells expressing their cognate ligand from a minigene (**Fig. 1D**). Each of these T cells, including the B7 TCR which recognizes the same target as the A6 TCR, also specifically killed T2 cells expressing their cognate antigens when co-cultured (**Fig. 1E**).

### PresentER minigene libraries can be used to discover the MHC-I peptide targets of T cells

In order to test the use of PresentER libraries to distinguish MHC-I targets of T cells from irrelevant targets, we harvested and activated splenocytes from OT-1 mice and wild type C57BL6/N mice. T cells from OT-1 mice express only H-2Kb/SIINFEKL specific TCRs whereas splenocytes from B6 mice are polyclonal. Splenocytes were co-cultured with RMA/S cells expressing a library of 5,000 randomly selected wild-type H-2Kb peptides as well as several control peptides. The abundance of each minigene after several days of co-culture was assayed by Illumina sequencing. A schematic of the experiment is presented in **Fig. 2A**. As expected, demonstrating the specificity of the system, cells expressing the peptide target of the OT-1 TCR — SIINFEKL—were depleted by more than 1-log while no other minigenes were depleted (**Fig. 2B)**.

**Fig 2:**
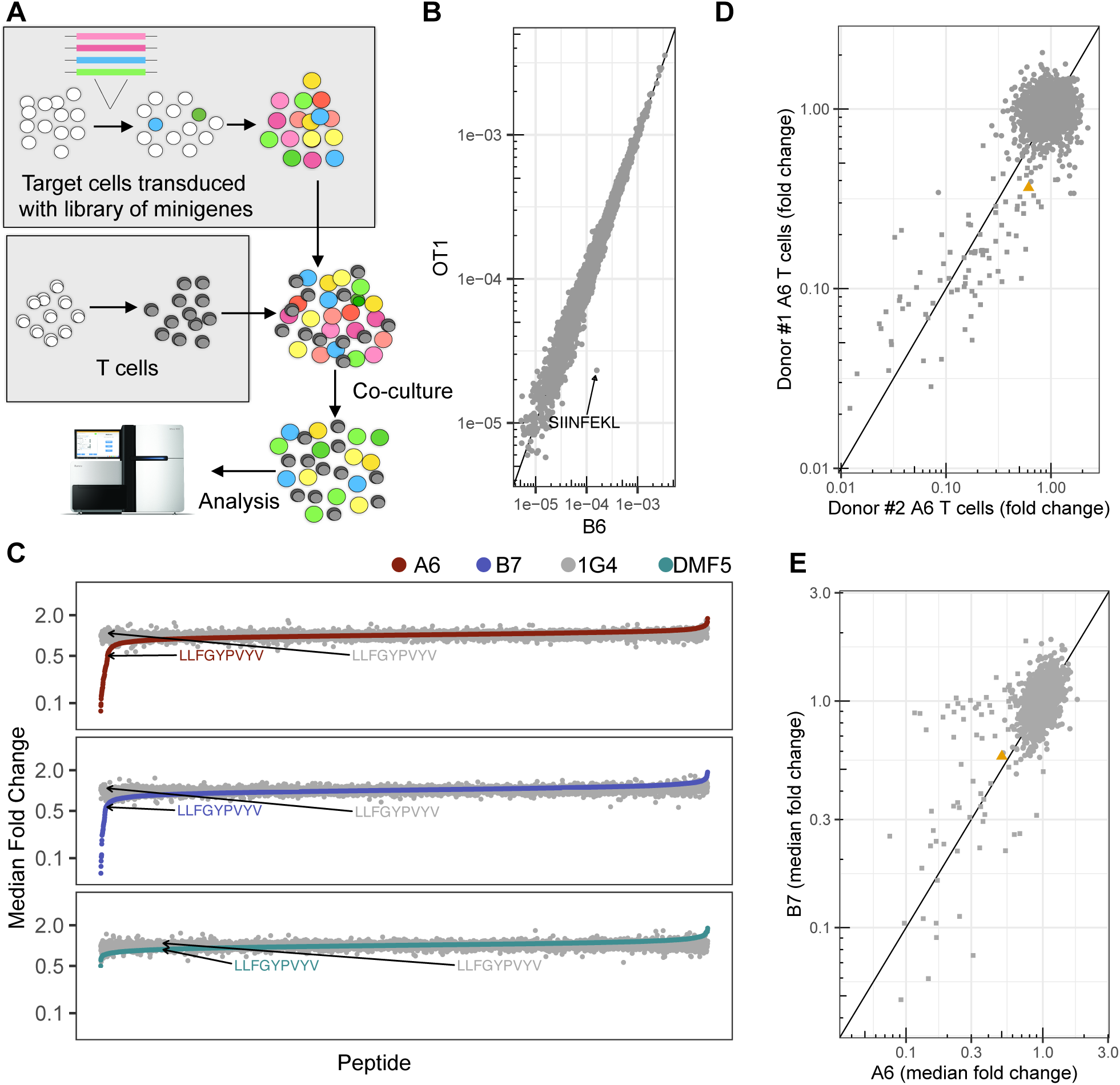
PresentER can be used to discover the peptide targets of T cell receptors. **(A)** A schematic depicting how cells expressing a library of minigenes are generated and subsequent co-cultured with native or engineered T cells. **(B)** The abundance of each minigene in a library of mouse H-2Kb antigens expressed by mouse RMA/S target cells after co-culture with activated OT-1 (y-axis) or B6 splenocytes (x-axis). **(C)** The median fold change of each peptide in the A6/B7 library after co-culture with human T cells from two different donors expressing the A6 TCR **(D)** The median fold change (across biological replicates) of each peptide in the library after co-culture with human T cells expressing the A6 (top), B7 (middle) or DMF5 (bottom) TCR, sorted on median fold change. LLFGYPVYV, the target of A6 and B7, is indicated. **(E)** The median fold change (across biological replicates) in abundance of each peptide in the A6/B7 library after co-culture with A6 (x-axis) or B7 (y-axis) expressing T cells

Next, we tested the ability of the PresentER system to discover the targets and off targets of two human TCRs. The A6 and B7 TCR were isolated from a patient with HTLV-1-associated myelopathy/tropical spastic paraparesis (HAM/TSP) and were found to recognize an HLA-A2.1 peptide derived from the HTLV-1 virus Tax protein. The A6 TCR in complex with its target was the first human TCR structure to be solved and the biochemical characteristics of both A6 and B7 have been extensively explored^26-28^. The position-specific binding specificities for the A6 and B7 TCR have been mapped by cytotoxicity studies using peptide-pulsed target cells^29^. These features made the A6 and B7 TCRs excellent candidates to use in a screen with the aim to validate previously known targets and discover additional targets. Using published data, we generated a motif of A6 and B7 binding and scored the human proteome for peptides that both matched the target peptide motifs and were predicted to bind to HLA-A2.1. We selected 5,000 peptides that were either highly scored for A6, B7 or both—as well as every single amino acid mutant to LLFGYPVYV—and cloned a library of minigenes expressing these peptides. T cells from non-HLA-A2.1 donors were engineered to express the A6, DMF5 or 1G4 TCR and co-cultured with the library of target cells. After co-culture, the abundance of each remaining minigene was quantified by Illumina sequencing. Fifty-two minigenes were depleted by 2-fold or more after co-culture with A6 T cells and 34 minigenes were depleted by co-culture with B7 T cells while no peptides were depleted by DMF5 or 1G4 (**Fig. 2C)**. To demonstrate the reliability of minigene depletion by T cell cytotoxicity, A6 T cell depletion by two T cell donors was compared. The minigenes that were depleted more than 2-fold between the two donors was 90% concordant (**Fig. 2D**). All the depleted minigenes were single amino acid substitutions of the Tax peptide. As expected based on prior data^29^, the A6 TCR is more promiscuous, recognizing almost twice as many peptides as B7 **(Fig. 2E)**.

In order to examine the ability of the PresentER screen to discover peptide targets of A6 and B7, we focused on the minigenes encoding single amino acid variants of the Tax peptide. These peptides had been evaluated for A6 and B7 cytotoxicity by Hausmann^29^ therefore we knew which were targets of the A6 and B7 TCRs. Using data from Hausmann, we constructed a position-specific scoring matrix (PSSM) for the peptide targets of A6 and B7 and scored every peptide (**Fig. 3A-B**). Most A6 and B7 ligands identified by Hausmann were depleted at least 2-fold in our minigene depletion assay **(Fig. 3C-D**). Using minigenes, we identified 19 and 17 respective Tax peptide substitutions that led to A6 and B7 cytotoxicity but were not tested by Hausmann; for instance substitutions at the 2^nd^ and 9^th^ position (revealing unusual MHC anchors), substitutions containing cysteines and differences between glutamic acid and aspartic acid residues in the same positions **(Fig. 3E-F)**.

**Fig. 3:**
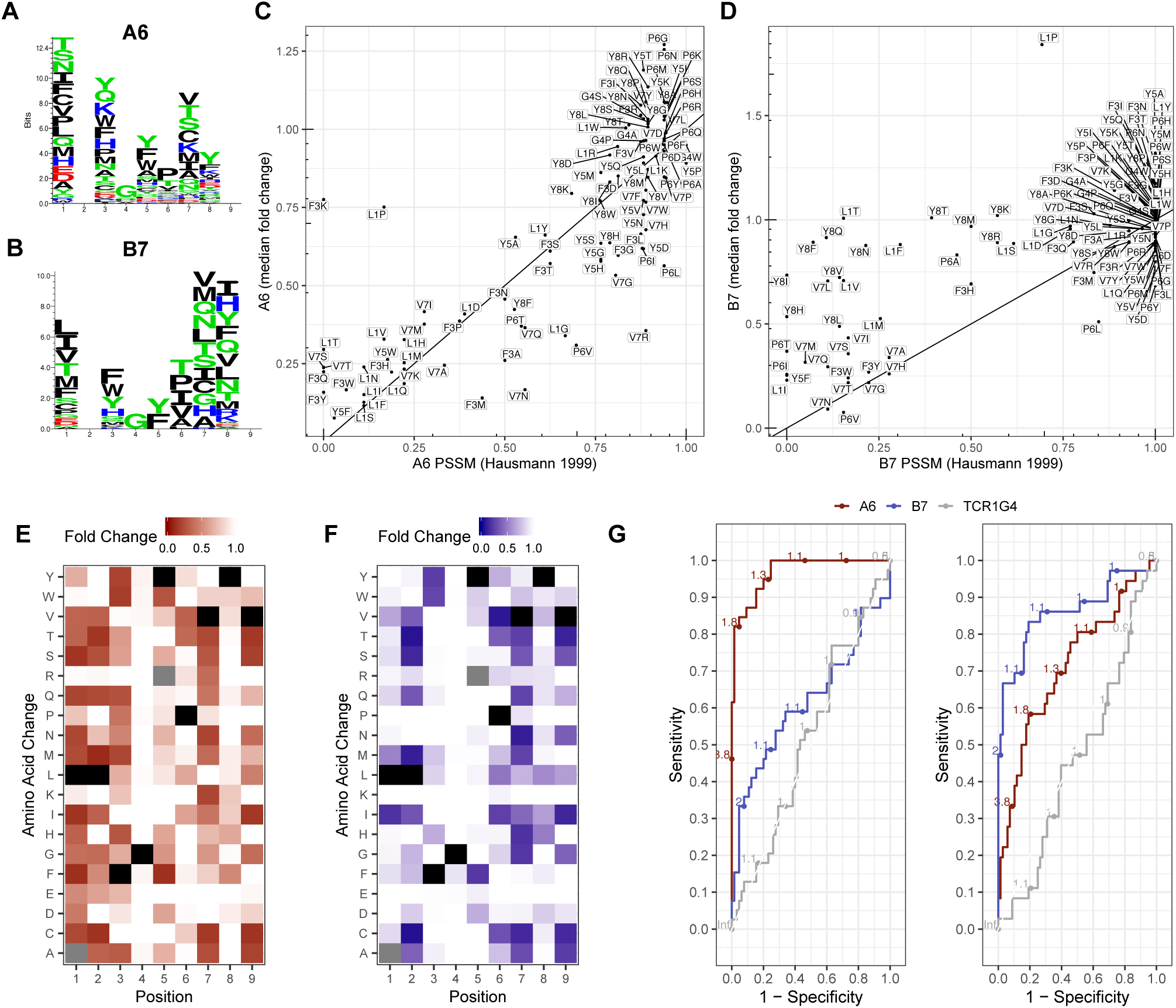
Previously identified single amino acid variants of the A6 and B7 target peptide are re-identified in the PresentER library screen. **(A-B)** Sequence logos showing the position specific frequency matrix (PSSM) of A6 and B7 derived from Hausmann et al (1999). **(C-D)** The abundance of each minigene encoding single amino acid mutants of the Tax peptide after co-culture with A6 or B7 (y-axis) in comparison to its PSSM score (x-axis). **(E-F)** Heat maps of the single amino acid substituted peptides obtained in the A6 and B7 minigene depletion assay experiments, respectively. **(G)** Receiver operating curves (ROC) showing that the sensitivity/specificity of the PresentER method using the A6 (left) and B7 (right) TCR experiments as compared to data from the literature on the binding specificity of these two TCRs.

A challenge that arises when evaluating new screening technologies is assessing its performance characteristics. In order to calculate sensitivity and specificity, it is necessary to know the true positives and negatives in a test population. In the case of T cell receptors, there is infrequently sufficient data to know the true and false positive rate. Moreover, the cost of empirically testing TCR reactivity for a meaningfully large dataset is prohibitive. The A6 and B7 TCR are unique in that a large number of peptides have been tested: 105 out of 133 possible single amino acid mutants of the LLFGYPVYV peptide were tested by Hausmann et al in co-culture killing assays and reported as percent-cytotoxicity. We categorized peptides with >25% of maximum killing in Hausmann’s assay as “positive” and the rest as “negative” and then calculated the performance characteristics of the PresentER A6 and B7 screens **(Fig 3G)**. The A6 TCR screen was highly sensitive, reaching >90% sensitivity with 85% specificity. The B7 TCR screen was less effective, reaching 70% sensitivity with 85% specificity. 1G4 is plotted for comparison. The sensitivity of PresentER is likely underestimated by this analysis, as the non-physiologic conditions of peptide pulsing *in vitro* tends to overestimate the T cell activating potential of a peptide.

Two peptides that had previously been characterized as weak off targets of the A6 and B7 TCRs in peptide-pulsing assays were also included in the library: S. Cerevisiae Tel1P 549–557 MLWGYLQYV and Human HuD / ELAVL4 87–95 LGYGFVNYI^29^. Neither of these peptides were depleted in the minigene library depletion assays, prompting us to wonder whether these peptides were indeed recognized well by the T cells. We performed an ELISPOT, using T2 cells pulsed with peptide or expressing PresentER minigenes. A6 was weakly reactive to the ELAVL4 peptide when pulsing onto T2 cells, but not to ELAVL4 minigene expressing cells, Tel1p peptide pulsed cells or Tel1p minigene expressing cells. B7 was not reactive to Tel1p or ELAVL4 by ELISPOT (**Fig. S6A**). T2 killing assays using minigene expressing peptides were also negative (**Fig. S6B**). The discrepancy between TCR recognition of peptide pulsed cells and minigene expressing cells indicates that the positive hits that emerge from PresentER minigene screening may be more reliable than hits which emerge from screens of peptide pulsed targets because peptide pulsed targets present peptide at supraphysiologic levels and thus may lead to false positives. We speculate that some of these peptides may not be well presented when expressed genetically, either because of inefficient loading onto MHC-I or removal during peptide editing, e.g. by TAPBPR^30^.

### PresentER minigene libraries can be used to discover the peptide targets of TCR mimic antibodies

The anti-cancer antibodies Pr20 and ESK1 that were developed by our group have been extensively evaluated to determine their sequence specificity. However, it was unknown whether their specificity was high enough to avoid cross-reacting with MHC-I peptides found on human cells. These antibodies were developed using phage display libraries and thus never underwent a thymic-like negative selection process. Based on alanine/residue scanning and structural^31^ data, we determined that ESK1 binding to RMFPNAPYL depended primarily on the R1 and P4 residues. Pr20 binding was mainly to the C-terminus of the peptide^22^. Therefore, we constructed a library of possible ESK1 and Pr20 cross-reactive targets by searching the human proteome *in silico* for 9 and 10-mer peptide sequences matching a motif based on prior biochemical data **(Fig. 4A)**. We located 1,157 and 24,539 potential cross-reactive peptides of ESK1 and Pr20, respectively, with NetMHCPan^32^ predicted HLA-A*02:01 affinity of less than 500nM. We synthesized a library of 12,472 oligonucleotides that together encoded all of the ESK1 cross reactive peptides and half of the Pr20 cross-reactive targets plus the single amino acid mutants of RMF and ALY (termed “CR-ESK1” and “CR-Pr20”, respectively), as well as positive/negative controls **(Fig. 4B)**. Library transduced T2 cells were stained with ESK1 or Pr20, sorted and sequenced. A schematic of the flow-based screen is presented in **Figure 4C**.

**Fig. 4:**
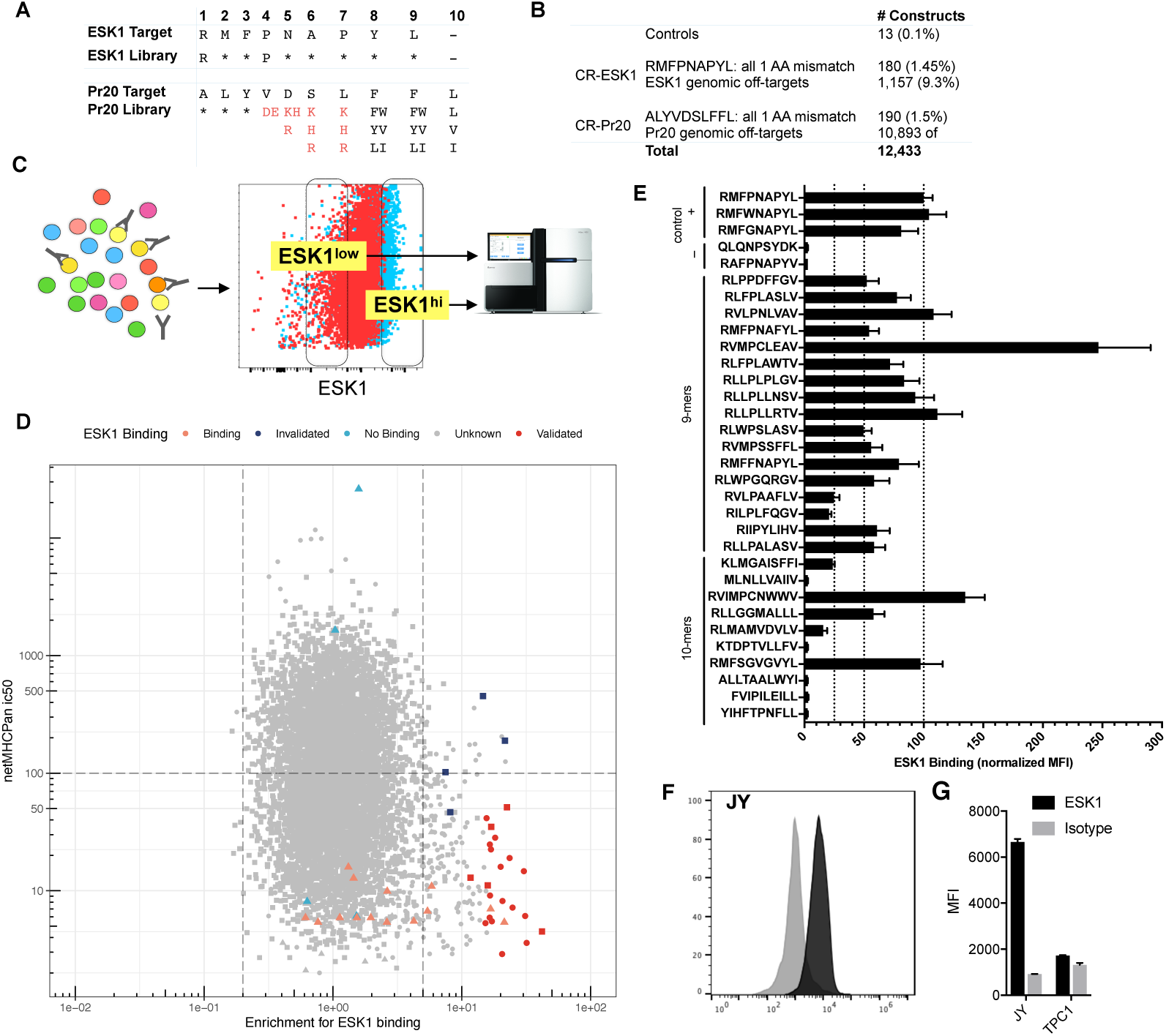
Discovery of off-targets of the ESK1 TCR mimic antibody. **(A)** The motif used to search the human proteome for peptides that might bind to ESK1 and Pr20. Asterisks indicate that any amino acid is allowed. Red characters indicate prohibited amino acids and black characters indicate allowed amino acids at that position. **(B)** Description of the constructed library. **(C)** Schematic of the flow-based screen. T2s are transduced at low MOI with retrovirus encoding a pool of PresentER minigenes, selected with puromycin and stained with the TCR mimic antibodies. Fluorescent activated cell sorting (FACS) is used to sort antibody binding and non-binding populations for sequencing. **(D)** Scatterplot of the ESK1 library screen. Each point is a unique peptide minigene plotted as minigene enrichment for ESK1 binding (x-axis; with 1 set as no enrichment) vs. predicted ic50 (in nM) to HLA-A*02:01 (y-axis). Lower ic50 indicates higher affinity. Control peptides and known ESK1 targets are plotted as triangles; CR-ESK1 as circles and CR-Pr20 as squares. Peptides that validated by peptide pulsing are displayed in dark red. Peptides that did not validate by peptide pulsing are in dark blue. **(E)** 27 peptides that were highly enriched for ESK1 binding and had high predicted affinity to HLA-A*02:01 (from Figure D) were synthesized at microgram scale, pulsed onto T2 cells and stained with a fluorescently labeled ESK1. Previously identified cross-reactive targets were included as positive controls. The median fluorescence intensity (MFI) of ESK1 binding is plotted, normalized to RMFPNAPYL (set at 100 units). **(F)** Representative ESK1 and isotype staining of the JY cell line. **(G)** Quantification of ESK1 and isotype staining of the JY and TPC1 cell lines.

Minigenes encoding previously known ESK1 ligands were enrichment for ESK1 binding compared to minigenes encoding non-ligands (p=0.032), suggesting that the flow-based screen was able to separate ESK1 binders from non-binders (**Fig. S7A)**. We found over 100 peptides with ic50 < 500nM and which were >5-fold enriched for ESK1 binding. Surprisingly, several of the most enriched peptides that emerged in the ESK1 screen were CR-Pr20 peptides, such as RVIMPCNWWV and RMFSGVGVYL (**Fig. 4D)**. Although these peptides are 10-mers, some bear sequence similarity to the target ligand of ESK1. To validate screen hits, we synthesized a subset of the enriched peptides and assayed them for binding to ESK1 by pulsing T2 cells. Of the 27 peptides tested, 22 (81%) showed increased binding to ESK1, including several which had originally been selected for Pr20 cross-reactivity and did not contain a proline in position 4 (**Fig. 4E)**. These unusual targets could not have been predicted from either the crystal structure of ESK1 or the alanine scanning data.

Next, we were interested to find out if any of the ESK1 targets we had discovered were expressed in a WT1-negative cell line, thus possibly leading to ESK1 binding. Recently, large databases of HLA-A*02:01 peptide ligands isolated from tumors and normal tissue have become available^15,33-35^. Within these databases (including personal correspondence with Department of Immunology members at Tübingen), we found two WT1-negative^36^ cell lines that contained ESK1 off-targets discovered in the PresentER screen (TPC-1: *RLPPPFPGL, RVMPSSFFL, RLGPVPPGL*, JY: *KLYNPENIYL, RLVPFLVEL*). RMFPNAPYL was not found among the MHC-I ligands immunoprecipitated from these lines. We tested ESK1 binding in each of these lines and found that JY cells bound ESK1 at high levels while TPC-1 was marginally positive for ESK1 binding (**Fig. 4F-G**). Thus, PresentER may be used to identify both theoretical and, in some cases, peptides presented on real cells that are bound by ESK1.

A screen of Pr20 cross-reactive ligands was performed in the same manner as described for ESK1. Positive control Pr20 binders were not enriched relative to the negative controls (p=0.71) (**Sup. Fig. 7B**), but twenty peptides were more than 5-fold enriched for Pr20 binding with predicted ic50s of less than 100nM (**Sup. Fig. 8**). We were able to synthesize and test 13 of these peptides by peptide pulsing and found that they were all validated as Pr20 binders.

### Peptide-MHC affinity influences the identified targets of TCR mimic antibodies

Examining only the CR-ESK1 subset of peptides, we noticed that the peptides most enriched for ESK1 binding were also predicted to have the highest affinity for HLA-A*02:01 (**Fig. 5A)**. The peptides that are ≥5-fold enriched for ESK1 binding have a higher affinity for HLA-A*02:01 as compared to the library as a whole or the peptides that were ≥5-fold depleted (median affinity of 31nM, 95nM and102nM, respectively) (**Fig. 5B**). We found the same result in the Pr20 screen: the most enriched Pr20 ligands also had the highest affinity to MHC-I. (**Fig. 5C**). The skew we observed in both ESK1 and Pr20-enriched minigenes towards high-affinity HLA-A*02:01 ligands suggests that genetic expression of peptides selects for presentation of ligands with the highest affinities for HLA-A*02:01. This may be an unexpected feature of PresentER, as affinity to MHC-I is the most important factor in determining if a peptide is presented on MHC-I (although high expression levels may overcome low affinity)^14^. Finally, we cloned minigenes for four of the most enriched CR-ESK1 (RLFPLAWTV 31.8x; KLMGAISFFI 41.9x) and CR-Pr20 (WLLGDSSFFL 6.5x; LLIQEGPFFV 6.6x) peptides and tested them for binding to ESK1 and Pr20. Cells expressing these peptides bound ESK1 and Pr20 at exceptionally high levels—significantly higher than the originally peptides to which ESK1 and Pr20 were designed (**Fig. 5D**).

**Fig. 5:**
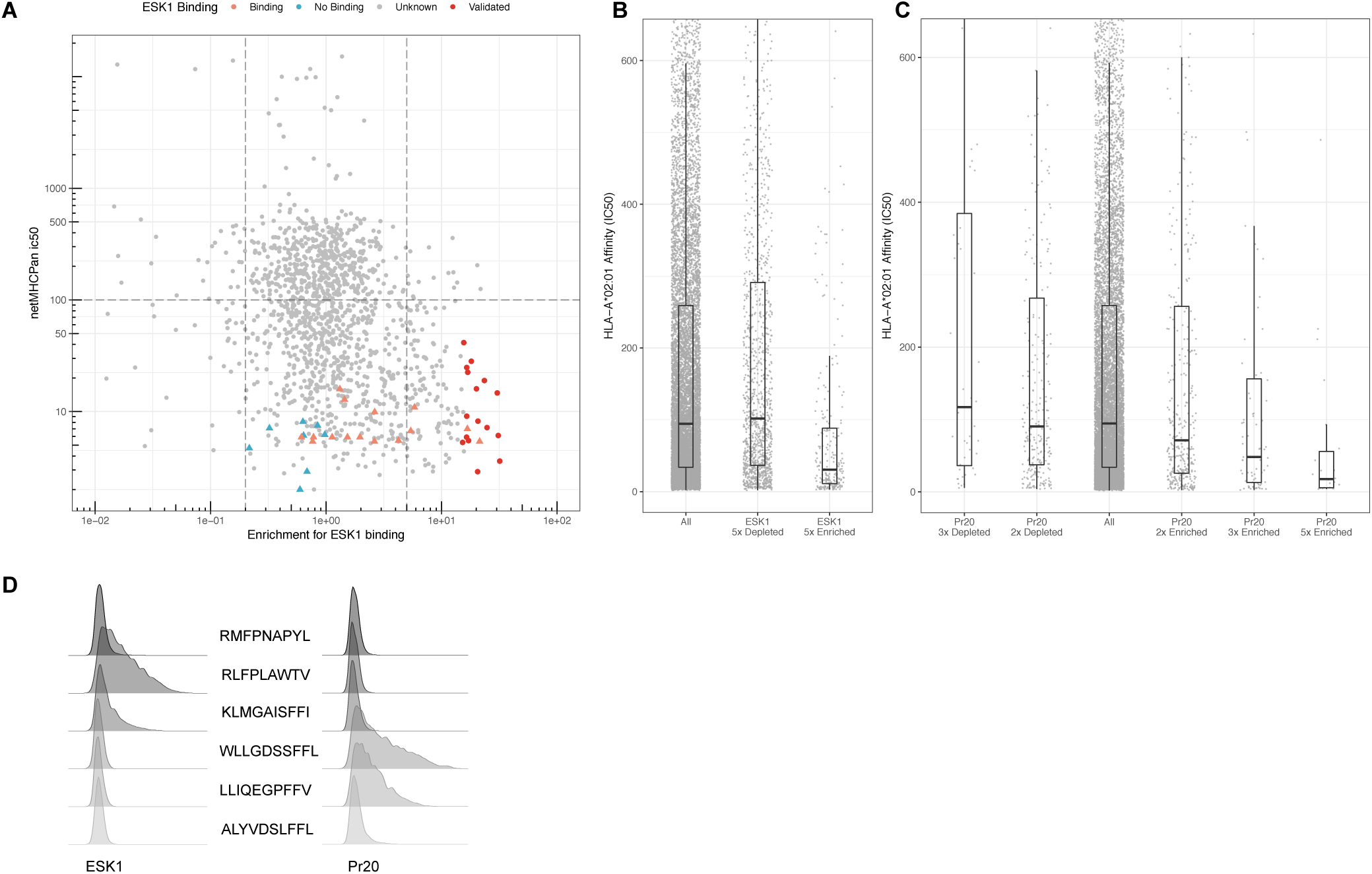
Peptides enriched in TCRm screening are high affinity MHC-I ligands. **(A)** Scatterplot of the ESK1 screen with only CR-ESK1 peptides and controls plotted. Each point is a peptide minigene plotted as enrichment for ESK1 binding (x-axis; 1 is no enrichment) vs. netMHCPan predicted HLA-A*02:01 affinity in ic50 (y-axis; nM). Lower ic50 indicates higher affinity. Control peptides and previously known ESK1 target peptides are plotted as triangles and CR-ESK1 as circles. Validated ESK1 binders are displayed in dark red. **(B)** The predicted HLA-A*02:01 affinity in ic50 (nM) of all screened peptides compared to peptides which were ≥5-fold depleted or ≥5-fold enriched for ESK1 binding. **(C)** The predicted HLA-A*02:01 affinity in ic50 (nM) of all screened peptides compared to peptides that were ≥5-fold, ≥3-fold or ≥2-fold enriched in the Pr20 screen and peptides that were ≤3 or ≤2-fold depleted for Pr20 binding. **(D)** ESK1 and Pr20 staining of T2 cells expressing 6 PresentER minigenes that were highly enriched in the screen.

## Discussion

Robust identification of the targets of TILs and the off-targets of TCR-based therapeutic agents and cells is not currently possible. As a consequence, preclinical evaluation of novel therapeutic agents directed towards peptide-MHC have been insufficient to prevent harmful off-tumor off-target toxicities, including deaths. A number of approaches have been developed that can identify the off targets of tumors, but all suffer from key limitations. For instance, animal models of cross-reactivity are not very useful due to species-specific MHC molecules and differences in peptide processing^37^. Yeast/Insect display is highly scalable, but relies on purified and refolded TCRs *in vitro*, which is not the native format that would be delivered to patients. Screening using MHC tetramers can be effective, but synthesis of large numbers of peptides is expensive. Methods to generate DNA-barcoded tetramers using *in vitro* transcription and translation are promising and we look forward to future development in this area. However, while these systems are valuable and have been used to elucidate the fundamental biology of TCRs^38^, they are poorly suited to preclinical evaluation of novel therapeutics. For instance, none of the methods described in the literature can be used to screen for off targets using T cell cytotoxicity assays, which is the least artificial assay that can be performed *in vitro*.

In PresentER, we have developed a mammalian approach to identifying MHC-I ligands of TCR-like agents at large scale. The PresentER system, by relying on the endogenous MHC presentation pathway of mammalian cells is a scalable, physiologically relevant system that can be used to identify functional cross-reactivities between MHC-I ligands and TCR agents with excellent sensitivity and specificity. In this report we have demonstrated that it can also be used for *in vitro* immunological assays while in a separate publication we demonstrate its use for *in vivo* immunological assays^10^. Minigenes covering all possible 9-10 amino acid peptides in the human exome that bind to one allele of MHC-I (<1×10^6^) could be constructed for $50,000, compared to the many months to years of work and the millions of dollars needed to do the same with synthesized peptides. This is a trivial sum for the preclinical evaluation of a novel engineered T cell therapeutic compared to the cost of a clinical trial that is halted because of toxicity^8,9^. Moreover, since PresentER relies on mammalian cells to display peptides—instead of covalently linking peptides directly to MHC on yeast—the antigen processing machinery of cells acts to prevent some peptides from reaching the surface, thus usefully limiting false positives. As an example of this, we have shown that peptides with weak MHC-I affinity are infrequently found to be off-targets of ESK1 and Pr20. A caveat to this approach is that a large number of target cells are required to ensure library fidelity—at least 1,000 cells per minigene—thus practically limiting the size of each minigene library.

In this report we have demonstrated that PresentER can be used for biochemical evaluation of therapeutic TCR based agents such as engineered TCRs and TCR mimic antibodies and discovered dozens of off targets of each. We believe that the minigene system we have described meets an unmet need in the preclinical evaluation of TCR agents. Given that patients have already died as a result of off-tumor off-target toxicity, we propose that off target assessment using libraries of MHC minigenes covering the entire human exome be a routine step in preclinical development of TCR and TCR-like agents.

## Acknowledgements

We thank Eureka Therapeutics for their generous gift of the ESK1 and Pr20 antibodies; Brian Baker and Lance Hellman for the gift of A6 and B7 TCR plasmids and Yael David for assistance in expressing and refolding; Scott Lowe and Eusebio Manchado Robles for the gift of the MSCV backbone and assistance with pooled library cloning; Steven A. Rosenberg for the DMF5 and 1G4 TCR. We thank the Integrated Genomics Operation at Memorial Sloan Kettering Cancer Center and the Flow Cytometry core facility for their assistance.

