## Supplemental Information for "Identification of the targets of T cell receptor therapeutic agents and cells by use of a high throughput genetic platform"

**Contents:**

1. **Supplementary Methods**
2. **Data availability**
3. **Table of DNA and Protein Sequences**
4. **Table of PresentER Constructs**
5. **Supplementary Figure 1 Overview of the PresentER Retroviral System.**
6. **Supplementary Figure 2 TCR and TCRm bind specifically to PresentER expressing cells.**
7. **Supplementary Figure 3 PresentER minigenes display peptides in TAP competent EL4 cells**
8. **Supplementary Figure 4 Mass spectrometry of PresentER peptide presentation and importance of signal peptide.**
9. **Supplementary Figure 5 PresentER is a single-copy competent expression system.**
10. **Supplementary Figure 6 Tel1p and ELAVL4 minigenes are not targets of A6 and B7**
11. **Supplementary Figure 7 Enrichment of ESK1 and Pr20 control peptides in library screens.**
12. **Supplementary Figure 8 Enrichment of Pr20 binding among library peptides**
13. **Supplementary References**

### Supplementary Methods

*Cloning PresentER Cassette and PresentER constructs*

The Mouse Stem Cell Virus vector that is the basis of the PresentER cassette was a generous gift from Scott Lowe of the Lowe lab. The 98 amino acid ENV_MMTVC signal peptide from Mouse Mammary Tumor Virus envelope protein (accession #Q85646) was found in the Signal Peptide Database (http://www.signalpeptide.de/index.php). IDT gBlocks encoding a modified, human codon optimized signal sequence followed by an antigen and a stop codon were ordered and cloned into the MLP vector with XhoI and EcoRI. The modifications changed two amino acids to enable directional cloning with the SfiI restriction enzyme: …`PQTSLTLFLALL`**`[S>A]`**`VL`**`[G>A]`**`PPPVSG`. A cassette with an SfiI site at the 3' end was also included in the gBlock and this construct is collectively termed the PresentER Cassette. In order to clone antigens into PresentER, DNA sequences encoding individual antigens were ordered from IDT, amplified with PresentER-F and PresentER-R primers and digested with SfiI. Digested inserts were purified with the Qiagen MinElute kit to remove primer dimers. The PresentER Cassette was digested with SfiI, treated with Calf Intestinal Phosphatase and gel purified. The inserts were ligated into the digested PresentER backbone with T4 ligase and transformed into NEB Stable cells. All sequences can be found in the table of DNA and protein sequences. The PresentER cassette and several example PresentER minigenes are available on Addgene.

*Production of retrovirus and library transduced cells*

HEK293T Phoenix amphoteric cells were transfected with polyethylenimine (PEI) and PresentER plasmid (15µg DNA : 45µg PEI) in 10cm TC plates. Virus was harvested every 12 hours, pooled, concentrated with Clontech Retro-X and frozen in aliquots. T2 cells were spinoculated with virus in non-TC treated 6-well plates at 32ºC x 2,000xg for 2 hours with 4µg/ml of polybrene. RMA-S cells were spinoculated with virus in non-TC treated 6-well plates at 32ºC x 1,000xg for 2 hours with 4µg/ml of polybrene. Library retrovirus was produced in the same way, except that virus production was scaled up to four 15cm plates. The volume of viral supernatant that led to 1/3 maximal transduction efficiency was established for each batch of virus produced. Transduced cells were selected with 1µg/ml (T2) or 4µg/ml (RMA/S) of puromycin for 2-3 days.

*Antibodies and commercial reagents*
ESK1 and Pr20 antibodies were purified by Eureka Therapeutics and labeled using Innova Biosciences lightning link kits (705-0010). Each labeled aliquot was titered on T2s or RMA-S cell bearing the PresentER minigenes. The TCR multimer specific to NLVPMVATV (CMV aa495-503)/HLA-A2.1 was purchased from Altor BioScience (Cat #TCR-CR1-0020). APC labeled antibodies specific to SIINFEKL/H2-Kb (Clone 25-D1.16) were purchased from Ebioscience (Cat #141606).

*Cloning the PresentER Library*
A pool of 12,472 oligonucleotides was synthesized by CustomArray, Inc in the following format: 5'-GGCCGTATTGGCCCCGCCACCTGTGAGCGGG…[27-30nt insert]…TAAGGCCAAACAGGCC-3'. The oligos were cloned into the PresentER vector in exactly the same way as the individual minigene. After ligation, the ligation products were electroporated into competent cells and plated onto four 15cm ampicillin plates. After overnight growth, the number of colonies was estimated to be ~$46 \times 10^6$. The colonies were scraped off the plate and grown for 3.5h in TB + ampicillin at 37ºC at 225rpm. The bacteria were maxiprepped and library representation was checked by Illumina sequencing.

*Library screening by FACS*
T2 cells transduced with library virus or single-minigene controls were stained with ESK1-APC or Pr20-APC. Sorting gates were set-up based on the fluorescence of the stained single-minigene control cells (RMF and ALY). Each sort was performed two times for each antibody. Sorted cells were frozen for genomic DNA extraction and Illumina sequencing. Bulk pre-sort cells were frozen as well.

*Genomic DNA Extraction and Library Sequencing*
Genomic DNA was extracted from bulk or sorted cells with the Gentra Puregene kit. Genomic DNA was amplified with barcoded primers and sequenced by the Integrated Genomics Operation core at MSKCC.

*Bioinformatics*
The peptides included in the PresentER library were found in Uniprot TrEMBL database of reviewed and unreviewed human protein sequences. Substrings of unique 9 and 10 amino acid sequences were collected and affinity to HLA-A*02:01 was calculated using NetMHCPan. Peptides with predicted ic50<500nM were compared to the ESK1 and Pr20 cross-reactivity motifs to determine which should be included in the library. All potentially cross-reactive ESK1 ligands and half of the potential Pr20 ligands were included in the library.

After Illumina sequencing, reads were mapped to the PresentER minigene library with Bowtie2. Reads that did not map to the minigenes in the library were discarded. Data analysis was performed in R. In order to identify minigenes encoding antibody ligands, we normalized the abundances in the sorted samples by the abundances in the unsorted samples and then divided the abundance of each minigene in the "antibody high" sample by the abundance of each minigene in the "antibody low" sample. Peptides with very low representation after DNA sequencing (1/50,000 or fewer than 10 reads) were excluded from analysis.

*Screen validation by peptide pulsing*
Spot synthesized crude peptides (PepTrack libraries) were ordered from JPT Peptide Technologies for validation of screen hits. Peptides were resuspended in DMSO followed by PBS and pulsed onto T2 cells. Pulsed cells were stained with ESK1 or Pr20 to evaluate binding of antibody to pMHC.

*Evaluation of ESK1 binding to JY and TPC1*
JY and TPC1 cells were stained with unlabeled ESK1 or IgG1 isotype control (Eureka Therapeutics Cat #ET901) followed by anti-human IgG1 APC antibody (BioLegend Cat #HP6017).

*Generation of AviTagged A6 TCR and A6 tetramers*
The A6 TCR was a generous gift from Brian Baker's lab. The beta chain plasmid was modified to encode a C-terminal AviTag biotinylation site (GLNDIFEAQKIEWHE). Gibson cloning was used to insert the site with the following two primers: F: 5'-gcagaaaattgaatggcatgaaTAAGCTTGAATTCCGATCCGG-3' R: 5'-gcttcaaaaatatcgttcaggccGTCTGCTCTACCCCAGGC-3'. Both the alpha and beta chains were expressed in BL21 (DE3) bacteria and induced with 1mM IPTG. The beta chain was co-expressed with a plasmid encoding the BirA enzyme (Addgene #26624) and supplemental biotin (0.5mM D-Biotin). Inclusion bodies were harvested and the individual chains were purified and re-folded together according to previously described protocols[1,2].

*Generation of A6 and B7 TCR mammalian expression plasmids*
In order to generate full-length mammalian TCR sequences, we used Gibson assembly to clone the A6 and B7 alpha/beta chains into the pMSGV1 vector backbone. The alpha and beta chain were separated using a P2A site. The complete A6 and B7 TCR sequences can be found in the table of DNA sequences below.

*Generation of T cells with transgenic TCRs*
Plasmids encoding the DMF5 and 1G4 TCRs were provided as a kind gift from Dr. Steven A. Rosenberg. TCR retroviral transduction was performed as described previously[3]. Briefly, retroviral particles were generated by transient transfection of the retroviral packaging cell line 293GP cells with the pMSGV1-TCR plasmids and pRD114 plasmid using Lipofectamine 2000 (Life Technologies). Retroviral supernatant was harvested 2 days later, and used to transduce PBMC that were stimulated with soluble 50 ng/ml anti-CD3 (OKT3, Miltenyi Biotec) and 300 IU/ml rhIL-2 (Chiron) for 2 days prior to retroviral transduction. Retroviral transductions were performed on Retronectin (Takara) coated non-tissue culture treated 24-wells plates by spinoculation of the retrovirus at 2,000× g, 32°C for 2 hours, followed by addition of activated T cells to the retrovirus containing plates. Following overnight incubation at 37°C, 5% CO2, T cells were transferred to a tissue-culture treated 24-wells plate and expanded in T cell media supplemented with 300 IU/ml rhIL-2 (Chiron). Transduced T cells were used at 10-15 days post-transduction or cryopreserved until used in assays.

*ELISPOT and co-culture killing assays*
IFN-gamma release ELISPOTs were performed in 200µl of RPMI supplemented with 5% FBS. 50,000 F5 transduced T cells were incubated at 1:1 effector:target (E:T) ratios with T2 cells expressing PresentER minigenes. Controls included: no targets, wild type T2s, 50µg/ml peptide pulsed T2s and PHA treated cells. Co-culture assays was performed in a similar manner: 50,000 T cells were incubated per well of a U-bottom plate with mixtures of 50,000 T2 cells expressing PresentER minigenes in mCherry or GFP flavors. The percentage of GFP positive cells was compared to the percentage of mCherry cells to evaluate specific lysis. Control wells did not contain T cells. Samples were evaluated by flow cytometry at 21 and 45 hours after co-culture.

*Co-culture library depletion assays*
Mouse RMA/S cells were transduced at low MOI (<0.3) with a library of 5,000 PresentER minigenes encoding wild type H-2Kb peptides (NetMHCPan v4.0 predicted ic50 < 500nM) selected randomly from the mouse proteome (UniProt database UP000000589 of canonical mouse protein sequences). Transduced cells were selected with 4µg of puromycin and then co-cultured with activated OT-1 or activated wild type B6 splenocytes for 4 days in 96-well round bottom plates. Cells were pooled together daily, media was added and cells were re-aliquoted onto the plates to prevent media exhaustion and avoid well-specific effects. Finally, cells were harvested, genomic DNA was extracted and minigenes were quantified by Illumina sequencing.

The human T2 library depletion assays were performed in a similar fashion. T2 cells transduced with a library of HLA-A2.1 peptides and co-cultured with A6, B7, DMF5 or 1G4 T cells for 4 days. Cells were pooled together daily, media was added as needed and cells were re-aliquoted onto the plates to prevent media exhaustion and avoid well-specific effects. Finally, cells were harvested, genomic DNA was extracted and minigenes were quantified by Illumina sequencing. Minigenes with low abundance (frequencies of less than 1 in 50,000 reads) were not considered in the depletion analysis.

**Data Availability**
The minigene counts and enrichment scores for each ESK1 and Pr20 library screen are available in the supplementary materials. The PresentER library sequencing reads are available at the following DOIs:

| Description | DOI |
|---|---|
| Minigene sequencing of sorted T2 cells expressing a library of HLA-A*02:01 exomic peptides. Cells are sorted for high and low binding of the TCR mimic antibody ESK1. | DOI:10.5281/zenodo.1313110 |
| Minigene sequencing of sorted T2 cells expressing a library of HLA-A*02:01 exomic peptides. Cells are sorted for high and low binding of the TCR mimic antibody Pr20. | DOI:10.5281/zenodo.1326544 |
| Minigene sequencing of T2 cells expressing a library of off targets (derived from A6 and B7 binding motifs in Hausmann 1999) after co-culture with A6, DMF5 or 1G4 expressing T cells. | DOI:10.5281/zenodo.1341943 |
| Minigene sequencing of T2 cells expressing a library of off targets (derived from A6 and B7 binding motifs in Hausmann 1999) are co-cultured with A6, B7, DMF5 or 1G4 expressing T cells. | DOI:10.5281/zenodo.1342624 |
| Minigene sequencing of RMA/S cells expressing a library of wild-type mouse H-2Kb peptides (plus controls) after co-culture with activated B6 or OT-1 splenocytes. | DOI:10.5281/zenodo.1419780 |

**Table of DNA and Protein sequences**

| Name | Description | DNA (5'->3') | Protein |
|---|---|---|---|
| ENV_MMTVC | MMTV ENV signal sequence (wild type) | ATGCCTAATCATCAGTCCGGGTCACCTACCGGCA<br>GTTCAGACCTGCTCCTTGATGGCAAGAAACAACG<br>AGCCCATCTGGCGCTGAGGAGAAAACGGCGACGG<br>GAAATGCGCAAGATTAACCGAAAGGTGAGAAGAA<br>TGAATCTCGCACCCATTAAAGAAAAAACAGCCTG<br>GCAGCACCTGCAAGCTCTGATCTTCGAGGCGGAA<br>GAAGTGTTGAAGACTTCTCAAACTCCACAGACCT<br>CCCTCACGCTGTTTCTGGCACTCTTGTCTGTACT<br>CGGGCCCCCACCTGTGAGCGGG | MPNHQSGSPTGSS<br>DLLLDGKKQRAHL<br>ALRRKRRREMRKI<br>NRKVRRMNLAPIK<br>EKTAWQHLQALIF<br>EAEEVLKTSQTPQ<br>TSLTLFLALLSVL<br>GPPPVSG |
| PresentER signal sequence | PresentER signal sequence based on ENV_MMTVC with SfiI sites FLALL[S>A]VL[G>A]PP P | ATGCCTAATCATCAGTCCGGGTCACCTACCGGCA<br>GTTCAGACCTGCTCCTTGATGGCAAGAAACAACG<br>AGCCCATCTGGCGCTGAGGAGAAAACGGCGACGG<br>GAAATGCGCAAGATTAACCGAAAGGTGAGAAGAA<br>TGAATCTCGCACCCATTAAAGAAAAAACAGCCTG<br>GCAGCACCTGCAAGCTCTGATCTTCGAGGCGGAA<br>GAAGTGTTGAAGACTTCTCAAACTCCACAGACCT<br>CCCTCACGCTGTTTCTAGCACTCTTGGCCGTATT<br>GGCCCCGCCACCTGTGAGCGGG | MPNHQSGSPTGSS<br>DLLLDGKKQRAHL<br>ALRRKRRREMRKI<br>NRKVRRMNLAPIK<br>EKTAWQHLQALIF<br>EAEEVLKTSQTPQ<br>TSLTLFLALLAVL<br>APPPVSG |
| PresentER Signal sequence scramble #1 | | **ATGGAAGGGGCCAGAAGCCAACTGTTGCTGAATC<br>ACATCTTGACTCCTATGATTCTTAGACTGCGAGC<br>ACAACTAATTAGGGCGTGGCGCCCACATCCTAAC<br>CAGGATACTTCCACGAAAGGGAAACTGGTGCAGC<br>TCTTTAAGGCCACCAAGATGCCAAAACTCCATCG<br>AAAGAAGGAGAGACCTTCATTGGAAAGTGCGGGC<br>TCTTCACAAGTGACACCGGTAGCCGCTTCCACCC<br>TCCTCCGACTGGTGAAACGGCCCGAGTTCAATGG<br>CCGGGCAGCCGAACAGGACCTG** | **MEGARSQLLLNHI<br>LTPMILRLRAQLI<br>RAWRPHPNQDTST<br>KGKLVQLFKATKM<br>PKLHRKKERPSLE<br>SAGSSQVTPVAAS<br>TLLRLVKRPEFNG<br>RAAEQDL** |
| PresentER Signal sequence scramble #2 | | **ATGCAGCTCTTGACTCCTACTAAGACAAAAAACG<br>AATCAGCCATGGCTGCCGCGAAGGTGTCAGCAAA<br>GTCTCGACTCTTGGCAAGAAGACTGACCGTACTC<br>AAGTTCCGGCCAAGCCTGGACCTGGAACATAGTG<br>TGGCCATTCGCCCTTCCAAACAACGATCCCAAAT<br>GCAGACGCGGCTTGATCTGGGCAATCCTGGCCAA<br>GTGCAGAATCCACTAGGGGAAGGGATCATTTTTT<br>GGCACGAAGCGCGAAAATTGCTCAAAAGACCCCA<br>TACCCTGCTGGCCAGGGAGCCG** | **MQLLTPTKTKNES<br>AMAAAKVSAKSRL<br>LARRLTVLKFRPS<br>LDLEHSVAIRPSK<br>QRSQMQTRLDLGN<br>PGQVQNPLGEGII<br>FWHEARKLLKRPH<br>TLLAREP** |
| PresentER antigen oligo | The region encoding the antigen is denoted by Xs. Th construct is amplified with PresentER-F and PresentER-R before digestion. | GGCCGTATTGGCCCCGCCACCTGTGAGCGGGXXX<br>…XXXTAAGGCCAAACAGGCC | |
| PresentER-F | Forward primer for amplifying antigen oligo | CGACTCACTATAGGGCCGTATTGGCC | |
| PresentER-R | Reverse primer for amplifying antigen oligo | AGTGATTTCCGGCCTGTTTGGCC | |
| P5Primer | Forward primer for amplifying minigene for Illumina sequencing | AATGATACGGCGACCACCGAGATCT | |
| P7BarcodePrimer | Reverse, barcoded primer for amplifying minigene for Illumina sequencing. XXXXXX denotes the Illumina primer. | CAAGCAGAAGACGGCATACGAGATXXXXX<br>XGTGACTGGAGTTCAGACGTGTGCTCTTC<br>CGATC | |

| | | |
|---|---|---|
| A6 TCR mammalian expression vector alpha and beta chain separated by P2A site | ATGAAGTCTTTGCGCGTACTCTTGGTGATATTGTGGCTCCAATTGAG<br>TTGGGTGTGGTCCCAGCAGAAGGAAGTGGAGCAGAACTCTGGACCC<br>CTCAGTGTTCCAGAGGGAGCCATTGCCTCTCTCAACTGCACTTACAG<br>TGACCGAGGTTCCCAGTCCTTCTTCTGGTACAGACAATATTCTGGGA<br>AAAGCCCTGAGTTGATAATGTCCATATACTCCAATGGTGACAAAGAA<br>GATGGAAGGTTTACAGCACAGCTCAATAAAGCCAGCCAGTATGTTTC<br>TCTGCTCATCAGAGACTCCCAGCCCAGTGATTCAGCCACCTACCTCT<br>GTGCCGTTACAACTGACAGCTGGGGGAAATTGCAGTTTGGAGCAGG<br>GACCCAGGTTGTGGTCACCCCAGATATCCAGAACCCGGATCCTGCC<br>GTGTACCAGCTGAGAGACTCTAAATCCAGTGACAAGTCTGTCTGCCT<br>ATTCACCGATTTTGATTCTCAAACAAATGTGTCACAAAGTAAGGATTC<br>TGATGTGTATATCACAGACAAATGTGTGCTAGACATGAGGTCTATGG<br>ACTTCAAGAGCAACAGTGCTGTGGCCTGGAGCAACAAATCTGACTTT<br>GCATGTGCAAACGCCTTCAACAACAGCATTATTCCAGAAGACACCTT<br>CTTCCCCAGCCCAGAAAGTTCCTGTGATGTCAAGCTGGTCGAGAAAA<br>GCTTTGAAACAGATACGAACCTAAACTTTCAAAACCTGTCAGTGATTG<br>GGTTCCGAATCCTCCTCCTGAAAGTGGCCGGGTTTAATCTGCTCATG<br>ACGCTGCGGCTGTGGTCCAGCCGGGCCAAGCGGTCCGGATCCGGA<br>GCCACCAACTTCAGCCTGCTGAAGCAGGCCGGCGACGTGGAGGAG<br>AACCCCGGCCCCATGAGCATCGGCCTCCTGTGCTGTGCAGCCTTGT<br>CTCTCCTGTGGGCAGGTCCAGTGAATGCTGGTGTCACTCAGACCCC<br>AAAATTCCAGGTCCTGAAGACAGGACAGAGCATGACACTGCAGTGT<br>GCCCAGGATATGAACCATGAATACATGTCCTGGTATCGACAAGACCC<br>AGGCATGGGGCTGAGGCTGATTCATTACTCAGTTGGTGCTGGTATCA<br>CTGACCAAGGAGAAGTCCCCAATGGCTACAATGTCTCCAGATCAACC<br>ACAGAGGATTTCCCGCTCAGGCTGCTGTCGGCTGCTCCCTCCCAGA<br>CATCTGTGTACTTCTGTGCCAGCCGGCCAGGTCTTGCCGGGGGACG<br>ACCAGAGCAGTATTTCGGGCCAGGGACGCGCCTTACGGTAACAGAA<br>GACTTGAAGAATGTCTTTCCACCTGAGGTCGCCGTTTTTGAACCCTC<br>CGAGGCCGAAATAAGTCATACTCAAAAGGCGACTCTGGTGTGCCTC<br>GCCACCGGGTTTTACCCGGACCACGTAGAACTTAGCTGGTGGGTGA<br>ATGGTAAAGAGGTCCATAGCGGGGTGTGCACGGACCCACAGCCTCT<br>CAAGGAACAACCCGCTCTGAATGATTCCAGGTATTGTCTTAGCTCAC<br>GGCTTCGAGTGTCAGCTACTTTTTGGCAAGATCCCCGCAACCACTTC<br>CGCTGTCAAGTCCAGTTCTACGGGCTCTCGGAGAATGACGAGTGGA<br>CCCAGGATAGGGCCAAACCCGTCACCCAGATCGTCAGCGCCGAGG<br>CCTGGGGTAGAGCAGACTGTGGCTTCACCTCCGAGTCTTACCAGCA<br>AGGGGTCCTGTCTGCCACCATCCTCTATGAGATCTTGCTAGGGAAG<br>GCCACCTTGTATGCCGTGCTGGTCAGTGCCCTCGTGCTGATGGCTA<br>TGGTCAAGAGAAAGGATTCCAGAGGCTAG | MKSLRVLLVILWLQL<br>SWVWSQQKEVEQNSG<br>PLSVPEGAIASLNCT<br>YSDRGSQSFFWYRQY<br>SGKSPELIMSIYSNG<br>DKEDGRFTAQLNKAS<br>QYVSLLIRDSQPSDS<br>ATYLCAVTTDSWGKL<br>QFGAGTQVVVTPDIQ<br>NPDPAVYQLRDSKSS<br>DKSVCLFTDFDSQTN<br>VSQSKDSDVYITDKC<br>VLDMRSMDFKSNSAV<br>AWSNKSDFACANAFN<br>NSIIPEDTFFPSPES<br>SCDVKLVEKSFETDT<br>NLNFQNLSVIGFRIL<br>LLKVAGFNLLMTLRL<br>WSSRAKRSGSGATNF<br>SLLKQAGDVEENPGP<br>MSIGLLCCAALSLLW<br>AGPVNAGVTQTPKFQ<br>VLKTGQSMTLQCAQD<br>MNHEYMSWYRQDPGM<br>GLRLIHYSVGAGITD<br>QGEVPNGYNVSRSTT<br>EDFPLRLLSAAPSQT<br>SVYFCASRPGLAGGR<br>PEQYFGPGTRLTVTE<br>DLKNVFPPEVAVFEP<br>SEAEISHTQKATLVC<br>LATGFYPDHVELSWW<br>VNGKEVHSGVCTDPQ<br>PLKEQPALNDSRYCL<br>SSRLRVSATFWQDPR<br>NHFRCQVQFYGLSEN<br>DEWTQDRAKPVTQIV<br>SAEAWGRADCGFTSE<br>SYQQGVLSATILYEI<br>LLGKATLYAVLVSAL<br>VLMAMVKRKDSRG |
| B7 TCR mammalian expression vector alpha and beta chain separated by P2A site | ATGAAGTCTTTGCGCGTACTCTTGGTGATATTGTGGCTCCAATTGAG<br>TTGGGTGTGGTCCCAGCAACAGAAGAATGATGACCAGCAAGTTAAG<br>CAAAATTCACCATCCCTGAGCGTCCAGGAAGGAAGAATTTCTATTCT<br>GAACTGTGACTATACTAACAGCATGTTTGATTATTTCCTATGGTACAA<br>AAAATACCCTGCTGAAGGTCCTACATTCCTGATATCTATAAGTTCCAT<br>TAAGGATAAAAATGAAGATGGAAGATTCACTGTCTTCTTAAACAAAAG<br>TGCCAAGCACCTCTCTCTGCACATTGTGCCCTCCCAGCCTGGAGACT<br>CTGCAGTGTACTTCTGTGCAGCAATGGAGGGAGCCCAGAAGCTGGT<br>ATTTGGCCAAGGAACCAGGCTGACTATCAACCCAAATATCCAGAATC<br>CGGACCCGGCCGTTTATCAGCTGCGTGATTCAAAATCTTCGGATAAA<br>TCCGTGTGTCTGTTCACGGATTTTGATAGCCAGACCAATGTGTCCCA<br>GTCAAAAGATAGTGATGTGTATATTACCGATAAATGCGTTCTGGACAT<br>GCGCAGTATGGATTTCAAAAGCAATTCGGCCGTGGCGTGGTCAAATA<br>AATCGGATTTCGCATGTGCGAACGCGTTTAATAACAGCATCATCCCG<br>GAAGATACGTTCTTTCCGAGCCCGGAAAGCTCTTGTGATGTCAAGCT<br>GGTCGAGAAAAGCTTTGAAACAGATACGAACCTAAACTTTCAAAACC<br>TGTCAGTGATTGGGTTCCGAATCCTCCTCCTGAAAGTGGCCGGGTTT<br>AATCTGCTCATGACGCTGCGGCTGTGGTCCAGCCGGGCCAAGCGGT<br>CCGGATCCGGAGCCACCAACTTCAGCCTGCTGAAGCAGGCCGGCG<br>ACGTGGAGGAGAACCCCGGCCCCATGAGCATCGGCCTCCTGTGCT<br>GTGCAGCCTTGTCTCTCCTGTGGGCAGGTCCAGTGAATGCTGGTGT<br>CACTCAGACCCCAAAATTCCAGGTCCTGAAGACAGGACAGAGCATG | MKSLRVLLVILWLQL<br>SWVWSQQQKNDDQQV<br>KQNSPSLSVQEGRIS<br>ILNCDYTNSMFDYFL<br>WYKKYPAEGPTFLIS<br>ISSIKDKNEDGRFTV<br>FLNKSAKHLSLHIVP<br>SQPGDSAVYFCAAME<br>GAQKLVFGQGTRLTI<br>NPNIQNPDPAVYQLR<br>DSKSSDKSVCLFTDF<br>DSQTNVSQSKDSDVY<br>ITDKCVLDMRSMDFK<br>SNSAVAWSNKSDFAC<br>ANAFNNSIIPEDTFF<br>PSPESSCDVKLVEKS<br>FETDTNLNFQNLSVI<br>GFRILLLKVAGFNLL<br>MTLRLWSSRAKRSGS<br>GATNFSLLKQAGDVE<br>ENPGPMSIGLLCCAA<br>LSLLWAGPVNAGVTQ<br>TPKFQVLKTGQSMTL |

| | | |
|---|---|---|
| | ACACTGCAGTGTGCCCAGGATATGAACCATGAATACATGTCCTGGTA<br>TCGACAAGACCCAGGCATGGGGCTGAGGCTGATTCATTACTCAGTT<br>GGTGCTGGTATCACTGACCAAGGAGAAGTCCCCAATGGCTACAATG<br>TCTCCAGATCAACCACAGAGGATTTCCCGCTCAGGCTGCTGTCGGC<br>TGCTCCCTCCCAGACATCTGTGTACTTCTGTGCCAGCAGTTACCCTG<br>GAGGAGGCTTTTATGAGCAGTATTTCGGTCCTGGAACAAGGCTGAC<br>CGTGACGGAAGATTTGAAAAATGTCTTTCCCCCAGAGGTAGCAGTCT<br>TCGAGCCGTCCGAGGCCGAGATATCCCATACCCAGAAGGCAACCCT<br>TGTTTGCTTGGCAACGGGATTTTATCCAGATCATGTGGAATTGTCCT<br>GGTGGGTCAACGGCAAAGAGGTTCACAGCGGCGTCTGCACAGATCC<br>GCAACCACTCAAGGAACAGCCCGCTCTTAATGATTCTCGCTACTGTC<br>TGAGTTCCAGGTTGCGGGTCAGCGCTACTTTCTGGCAGGATCCCCG<br>CAACCACTTCCGCTGTCAAGTCCAGTTCTACGGGCTCTCGGAGAATG<br>ACGAGTGGACCCAGGATAGGGCCAAACCCGTCACCCAGATCGTCAG<br>CGCCGAGGCCTGGGGTAGAGCAGACTGTGGCTTCACCTCCGAGTCT<br>TACCAGCAAGGGGTCCTGTCTGCCACCATCCTCTATGAGATCTTGCT<br>AGGGAAGGCCACCTTGTATGCCGTGCTGGTCAGTGCCCTCGTGCTG<br>ATGGCTATGGTCAAGAGAAAGGATTCCAGAGGCTAG | QCAQDMNHEYMSWYR<br>QDPGMGLRLIHYSVG<br>AGITDQGEVPNGYNV<br>SRSTTEDFPLRLLSA<br>APSQTSVYFCASSYP<br>GGGFYEQYFGPGTRL<br>TVTEDLKNVFPPEVA<br>VFEPSEAEISHTQKA<br>TLVCLATGFYPDHVE<br>LSWWVNGKEVHSGVC<br>TDPQPLKEQPALNDS<br>RYCLSSRLRVSATFW<br>QDPRNHFRCQVQFYG<br>LSENDEWTQDRAKPV<br>TQIVSAEAWGRADCG<br>FTSESYQQGVLSATI<br>LYEILLGKATLYAVL<br>VSALVLMAMVKRKDS<br>RG |

**Table of PresentER Constructs**

| **Species / Protein / Position** | **DNA encoding peptide** | **Peptide** |
|---|---|---|
| Human WT1 126-134 | AGGATGTTTCCTAACGCGCCCTACCTG | RMFPNAPYL |
| Human PRAME 300-309 | GCTCTCTATGTGGACTCTTTATTTTTCCTT | ALYVDSLFFL |
| CMV pp65 495-503 | AACCTGGTGCCCATGGTGGCCACCGTG | NLVPMVATV |
| Human EW | CAGCTGCAGAACCCCAGCTACGACAAG | QLQNPSYDK |
| Influenza M peptide 58-66 | GGCATCCTGGGCTTCGTGTTCACCCTG | GILGFVFTL |
| Human WT1 239-247 | AACCAGATGAACCTGGGCGCCACCCTG | NQMNLGATL |
| Human MART-1 27-35 | GCGGCCGGAATAGGCATATTGACTGTA | AAGIGILTV |
| Human NY-ESO-1 157-165 | TCTCTGCTGATGTGGATCACTCAATGT | SLLMWITQC |
| HTLV-1 Tax 11-19 | CTGCTTTTTGGTTACCCTGTTTATGTT | LLFGYPVYV |
| Human ELAV-like protein 4 (HuD) 87–95 | CTTGGATATGGGTTTGTCAATTATATA | LGYGFVNYI |
| S. cerevisiae Tel1p 549-557 | ATGCTTTGGGGCTATCTGCAATACGTT | MLWGYLQYV |
| Human ADCY4 789-798 | AAACTCATGGGAGCAATCTCATTTTTTATA | KLMGAISFFI |
| Human EFNA1 186-194 | CGACTCTTTCCTCTCGCATGGACTGTC | RLFPLAWTV |
| Human SH3TC1 205-214 | CTCCTGATCCAAGAAGGGCCTTTCTTTGTT | LLIQEGPFFV |
| Human HARBI1 221-230 | TGGCTCCTCGGTGACTCTTCATTCTTCCTT | WLLGDSSFFL |
| Chicken ovalbumin 257-264 | AGCATCATCAACTTCGAGAAGCTG | SIINFEKL |
| Mouse PEDF 271-279 | ATGAGCATCATCTTCTTCCTGCCCCTG | MSIIFFLPL |

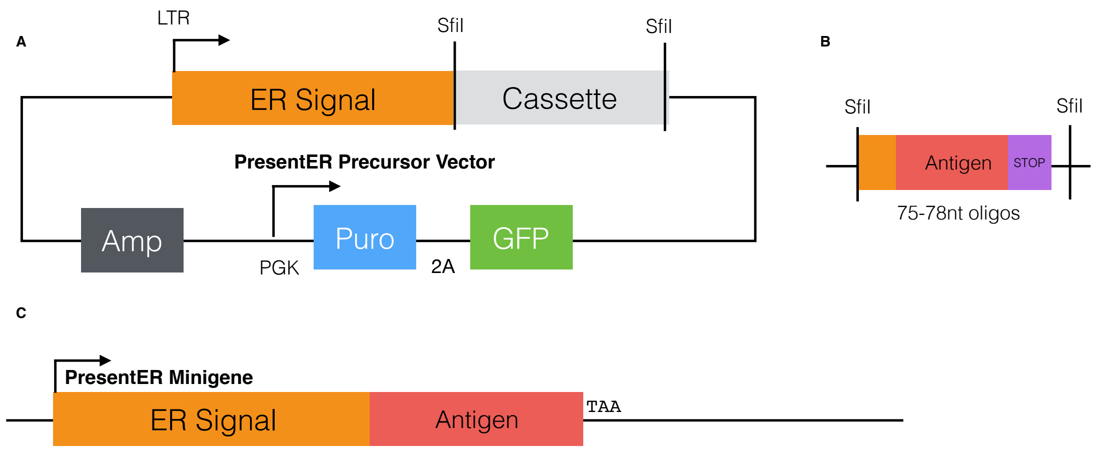


**Supplementary Figure 1:** Overview of the PresentER Retroviral System. **(A)** The PresentER system is based on an MSCV retroviral vector. The peptide antigen minigene is driven by the MSCV LTR and encodes an endoplasmic reticulum (ER) targeting sequencing followed by the precise peptide to be expressed, followed by a stop codon. The vector contains a puromycin resistance gene and GFP driven by PGK. The endoplasmic reticulum sequence is the leader sequence from MMTV gp70 protein. A removable cassette for easy cloning is bounded by SfiI restriction sites. **(B)** The oligonucleotide sequence encoding the antigen is an inexpensive 75-78nt sequence and can be amplified, digested with SfiI and ligated into the backbone. **(C)** The cloned PresentER minigene construct.

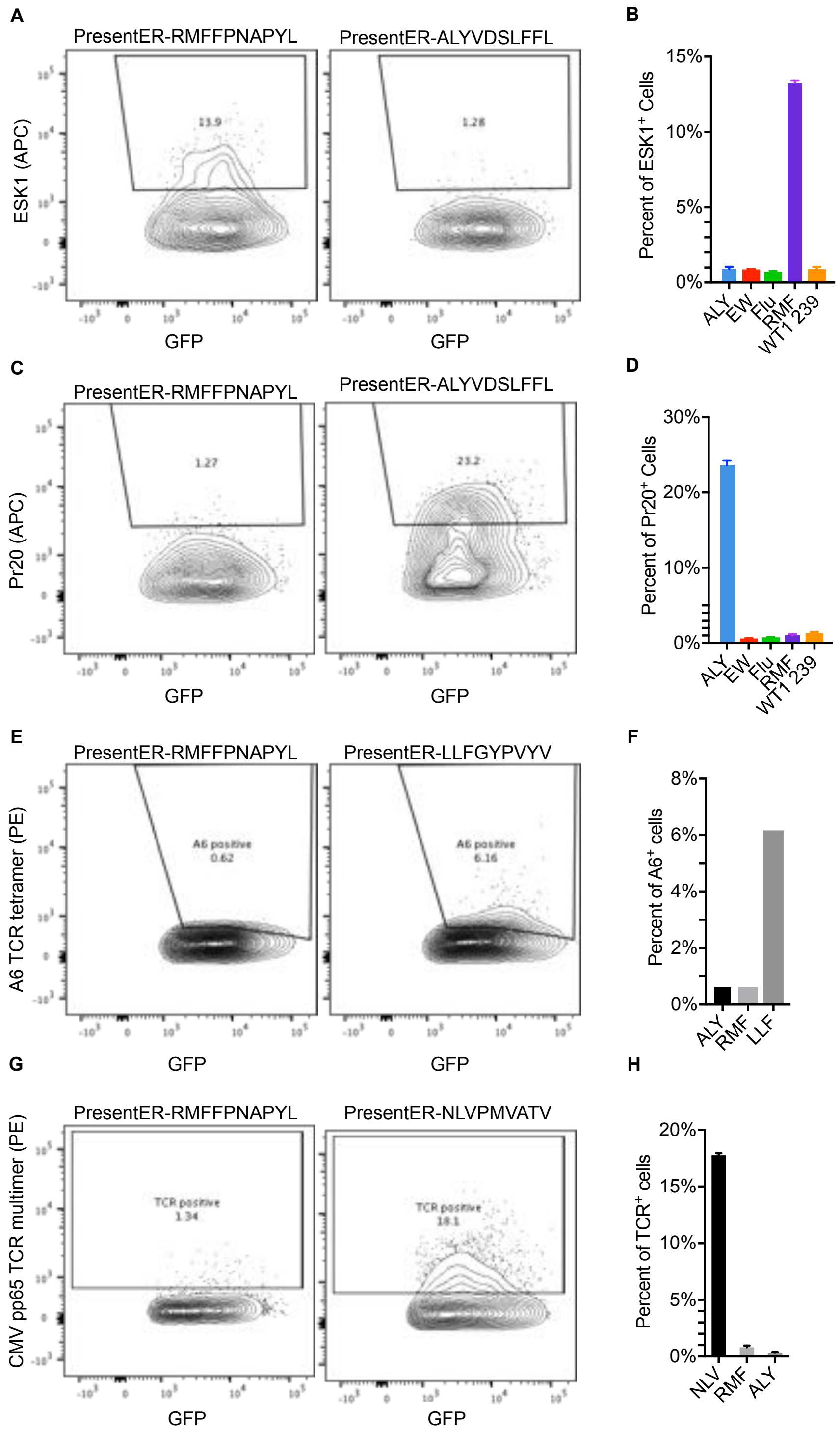


**Supplementary Figure 2:** TCR and TCRm bind specifically to PresentER expressing cells. Representative contour plots and quantification of the TCR mimic antibodies ESK1 **(A-B)** and Pr20 **(C-D)** binding to T2 cells expressing their PresentER targets of irrelevant controls. Representative contour plots and quantification of two TCR multimers, one specific to HTLV-1 Tax peptide (**E,F**) and one specific to CMV pp65 peptide (**G,H**) binding specifically to T2 cells expressing their target PresentER minigenes or irrelevant minigenes.

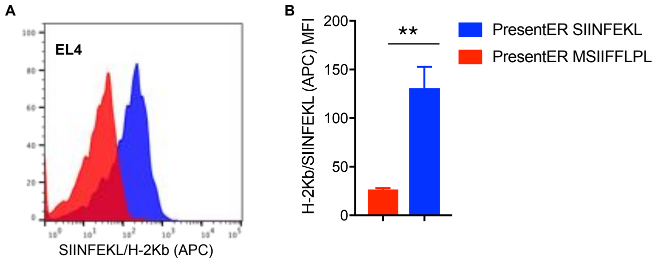


**Supplementary Figure 3:** PresentER minigenes display peptides in TAP competent EL4 cells Representative histogram **(A)** and quantification **(B)** of the binding of the SIINFEKL/H-2Kb specific antibody (25-D1.16) to EL4 cells expressing PresentER-SIINFEKL or a control H-2Kb peptide MSIIFFLPL.

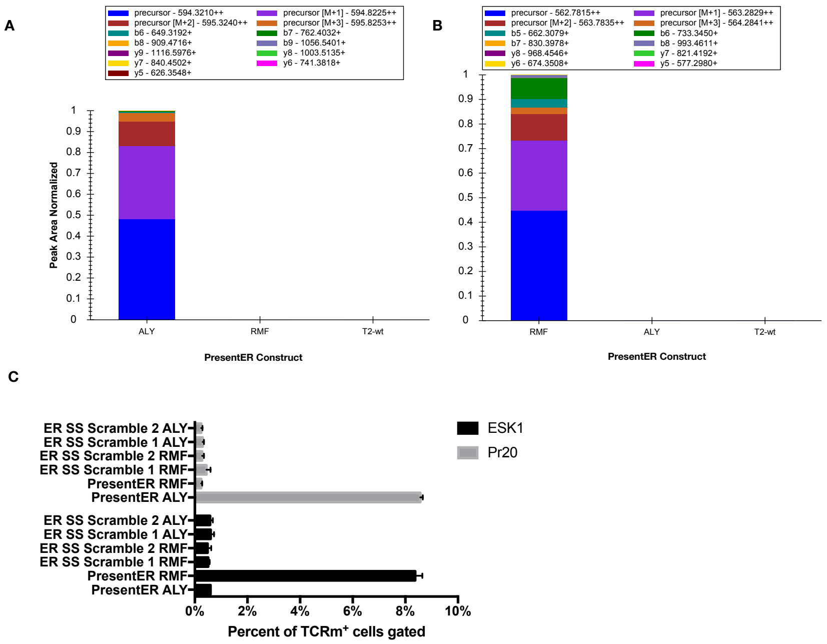


**Supplementary Figure 4:** Mass spectrometry of PresentER peptide presentation and importance of signal peptide. **(A-B)** MHC-I ligands were isolated from wild-type T2 cells or T2 cells expressing PresentER-ALY or PresentER-RMF. Peptides were identified by mass spectrometry and the intensity of ions corresponding to the ALY (**A**) or RMF (**B**) peptides in each sample are plotted. (**C**) ESK1 and Pr20 binding to T2 cells expressing PresentER-RMF and PresentER-ALY is compared with binding to T2 cells expressing RMF and ALY downstream of scrambled signal sequences.

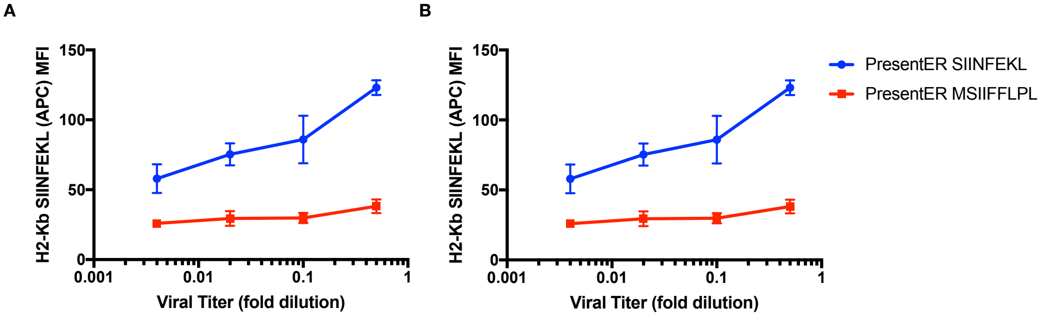


**Supplementary Figure 5:** PresentER is a single-copy competent expression system. RMA/S cells were transduced with serially diluted PresentER-SIINFEKL or PresentER-MSIIFFLPL virus in biological replicates. The percent of GFP positive, SIINFEKL/H-2Kb positive cells after transduction is plotted as a function of fold dilution of viral supernatant **(A)** and percent of cells infected **(B)**. Low infection efficiency does not lead to loss H2-Kb SIINFEKL binding, as would be expected for a minigene that was not single copy competent.

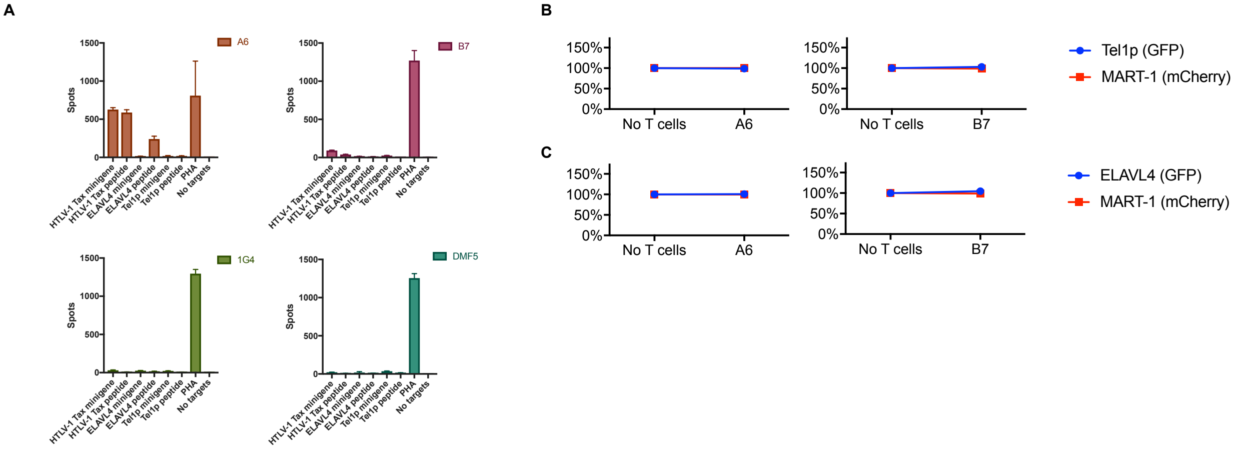


**Supplementary Figure 6: (A)** ELISpot of genetically engineered T cells expressing the A6 (target: HTLV-1 Tax 11-19 LLFGYPVYV), DMF5 (target: MART-1 27-35 AAGIGILTV) or 1G4 (target: NY-ESO-1 157-165 SLLMWITQC) TCRs challenged with peptide-pulsed T2 cells or T2 cells expressing PresentER minigenes. **(B)** Results of *in vitro* co-culture killing assays where A6 or B7 (reactive with Tax peptide) expressing T cells were incubated with a mixture of T2 cells expressing PresentER-Tel1p (GFP)/PresentER MART-1 (mCherry), PresentER ELAVL4 (GFP)/PresentER MART-1 (mCherry) or without any T cells. The change in abundance of the T2 target cells are plotted relative to their abundance in the "No T cells" sample.

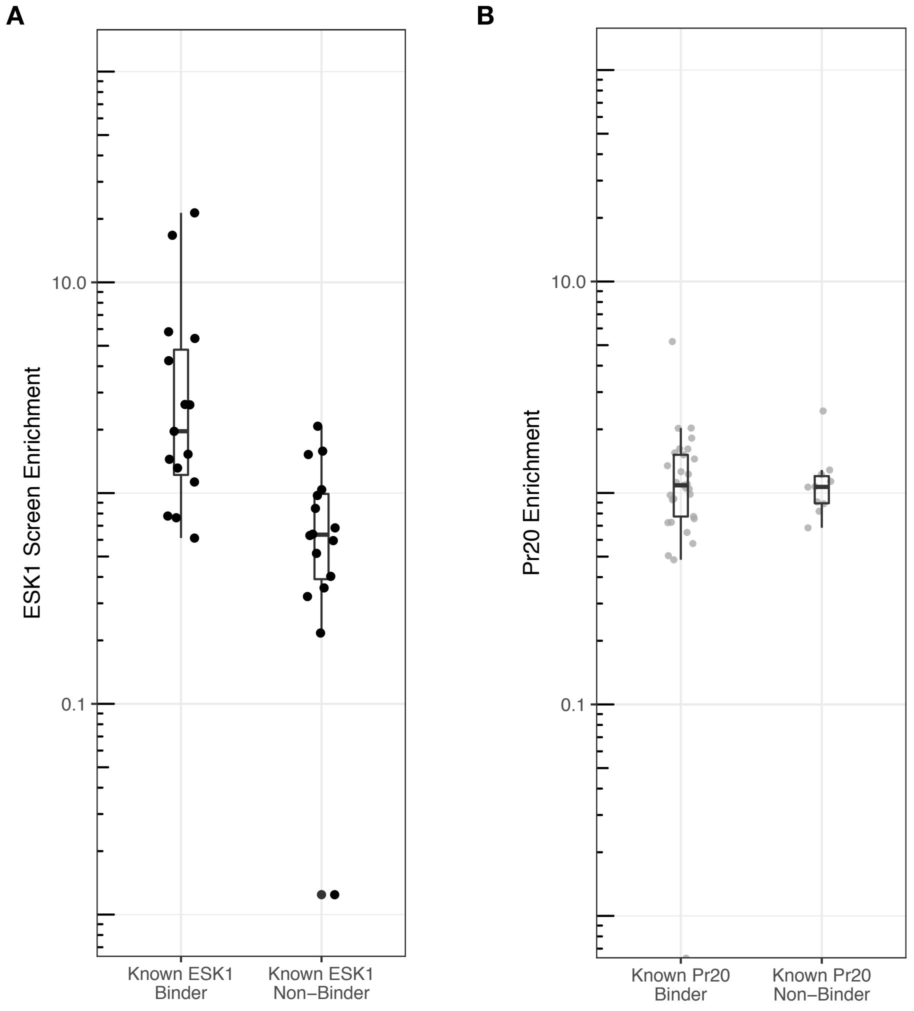


**Supplementary Figure 7:** Enrichment of ESK1 and Pr20 control peptides in library screens. **(A)** The flow screen enrichment scores of control peptides known to be ESK1 binders or non-binders. **(B)** The flow screen enrichment scores of control peptides known to be Pr20 binders or non-binders.

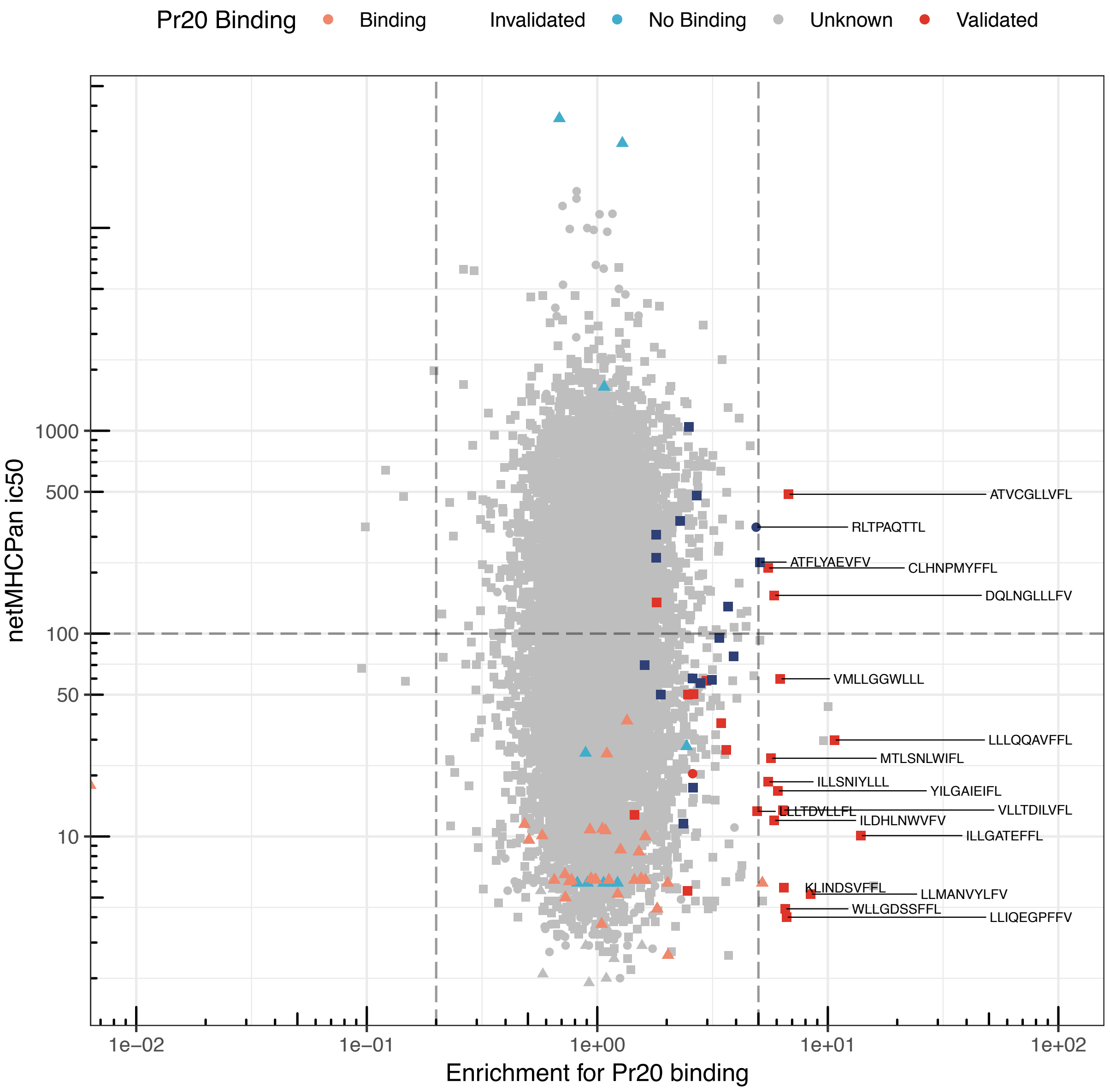


**Supplementary Figure 8:** Enrichment of Pr20 binding among library peptides. Each point is a unique peptide minigene with the x-axis indicating enrichment for Pr20 binding (with 1 set as no enrichment) and y-axis indicating the peptide's predicted ic50 (in nM) to HLA-A*02:01. Lower ic50 indicates higher affinity. Marked control peptides and known Pr20 peptide targets are plotted as triangles; CR-ESK1 peptides, as circles and CR-Pr20 peptides, as squares. Peptides that validated as Pr20 binders by peptide pulsing are displayed in dark red. Peptides that did not validate by peptide pulsing are in dark blue.
